# The *kpc-1* 3’UTR facilitates dendritic transport and translation of mRNAs for dendrite arborization of a mechanosensory neuron important for male courtship

**DOI:** 10.1101/2021.08.03.453128

**Authors:** Mushaine Shih, Yan Zou, Tarsis Ferreira, Nobuko Suzuki, Eunseo Kim, Chiou-Fen Chuang, Chieh Chang

## Abstract

A recently reported Schizophrenia-associated genetic variant in the 3’UTR of the human furin gene, a homolog of *C. elegans kpc-1*, highlights an important role of the furin 3’UTR in neuronal development(*1*). We isolate three *kpc-1* mutants that display abnormal dendrite arborization in PVD neurons and defective male mating behaviors. We show that the *kpc-1* 3’UTR participates in dendrite branching and self-avoidance. The *kpc-1* 3’UTR facilitates mRNA localization to branching points and contact points between sibling dendrites and promotes local protein synthesis. We identify a secondary structural motif in the *kpc-1* 3’UTR required for dendrite self-avoidance. Animals with *dma-1* receptor over-expression exhibit similar dendrite branching and self-avoidance defects that are suppressed with *kpc-1* over-expression. Our results support a model in which KPC-1 proteins are synthesized at branching points and contact points to locally down-regulate DMA-1 receptors to promote dendrite branching and self-avoidance of a mechanosensory neuron important for male courtship.

## Introduction

PVD nociceptive neurons innervate *C. elegans* skin with elaborate dendrite arbors that respond to harsh mechanical stimuli(*2*). Dendrite arborization allows uniform sensory coverage of PVD neurons across the entire surface of the animal outside the head region. However, the mechanism underlying this process is still largely unknown.

Dendrite arborization in PVD neurons is restricted within a defined time frame by two intrinsic timing mechanisms(*3*) and is constrained spatially by two organizing principles: Escape (branch out) from the intermediate target (the trap zone) upon arrival at it and self-avoidance(*4–6*). Self-avoidance is a process in which self-recognition among neurites of a single neuron is followed by their repulsion from each other. This selective repulsion between dendrites of the same cell was originally observed in leech neurons and proposed to promote a uniform receptive field(*7*, *8*). Recent studies using *Drosophila* as a model showed that homophilic binding of Dscam1 proteins on sister branches of the same cell (MB or da neurons) promotes repulsive interactions between them, to ensure that they diverge and grow along separate pathways(*9–13*). Since the Dscam1 mutation does not completely disrupt dendrite self-avoidance in *Drosophila* and there is no Dscam1 homolog in *C. elegans*, studying PVD dendrite self-avoidance in *C. elegans* provides a unique opportunity to identify novel mechanisms of dendrite self-avoidance. Indeed, several known developmental pathways, including UNC-6 (netrin), MIG-14 (Wntless), and FMI-1 (Flamingo) signaling, have been implicated in PVD dendrite self-avoidance in *C. elegans*(*6*, *14*, *15*).

It was previously reported that guidance cues SAX-7 (L1CAM) and MNR-1 (Menorin) generated in the hypodermis (skin) established dendrite growth pathways and instructed dendrite arborization for PVD neurons in *C. elegans*(*16*, *17*). PVD neurons first grow two longitudinally extending 1° dendrites along the anterior-posterior (A/P) axis. Orthogonal arrays of 2°, 3°, and 4° dendritic branches emerge sequentially in a manner that alternates between the dorsal-ventral (D/V) and the A/P axes to produce an elaborate network of sensory processes(*5*, *18*) (Fig. 1b). Skin cues, SAX-7 and MNR-1, and a muscle-derived cue, LECT-2, are required for dendrite pattern formation through direct interactions with the dendrite receptor DMA-1 to form a multi-protein receptor-ligand complex(*16*, *17*, *19–21*).

**Figure 1.**
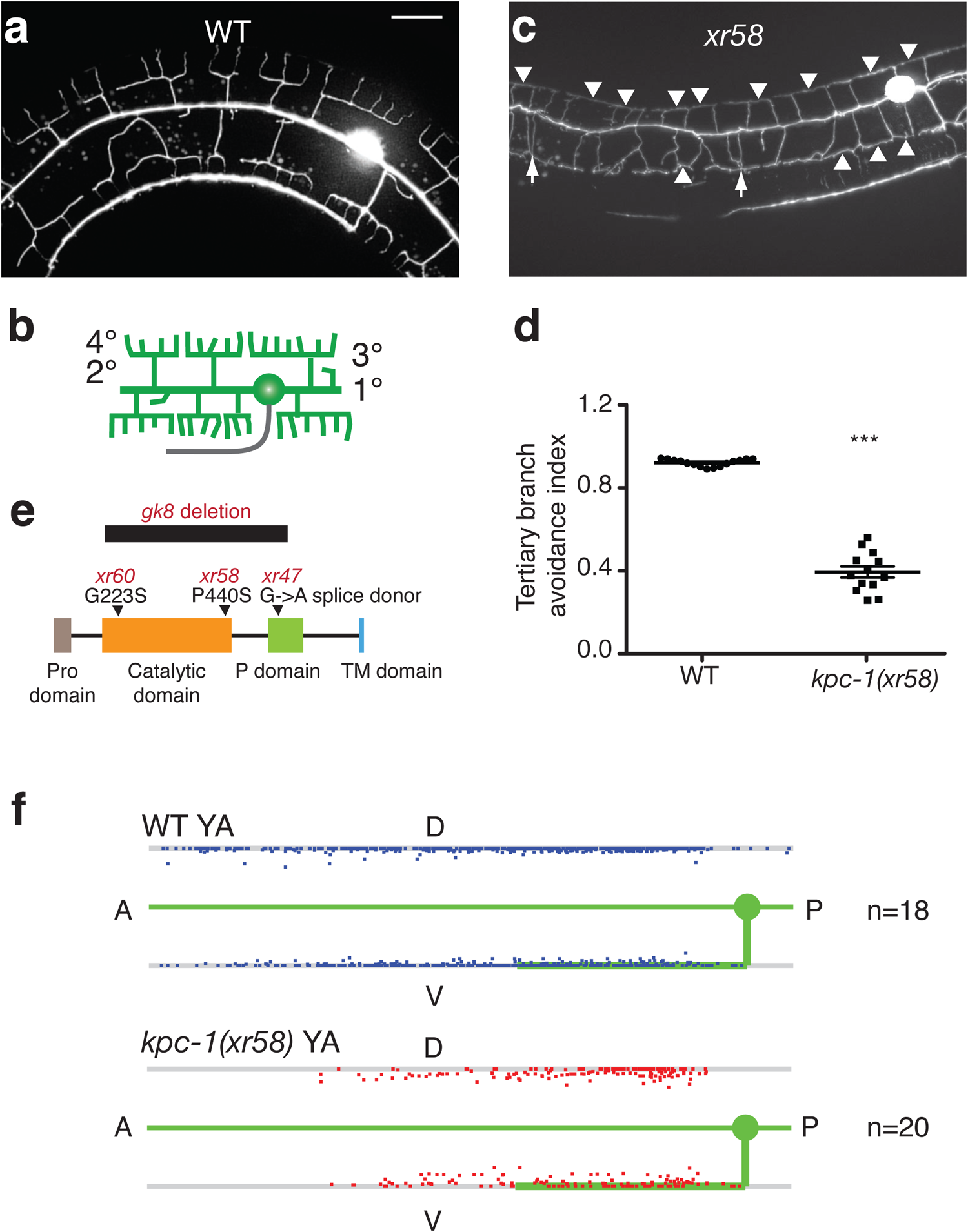
*kpc-1(xr58)* mutations preferentially affect PVD dendrite self-avoidance. (a, c) Images of PVD dendrites in wild type (a) and *xr58* mutants (c). Scale bar, 20 μm. (b) Schematic of dendrites (green) and axon (grey) extended from a PVD neuron. PVD projected primary (1°) dendritic processes anteriorly and posteriorly from the soma. Higher order dendritic branching (secondary 2°, tertiary 3°, quaternary 4°) alternated between anterior-posterior and dorsal-ventral directions to generate a regularly spaced array of parallel branches. *xr58* mutants, while showing effective secondary dendrite branching, displayed extensive defects in self-avoidance of sibling dendrites. Arrows point to contacts between neighboring secondary dendrites. Arrowheads point to contacts between neighboring tertiary dendrites. (d) Tertiary branch avoidance index (TBAI) in wild type versus *kpc-1(xr58)* mutants. *kpc-1(xr58)* mutants displayed significantly lower TBAI than wild-type animals. ***p < 0.001 by a Student’s *t*-test. (e) Schematic of the KPC-1 protein motifs showing locations of *xr47*, *xr58*, and *xr60* mutations. (f) Scatter plots showing the positions of 4° dendrite termini in wild-type young adults and *kpc-1(xr58)* young adults. In each scatter plot, the top line (dorsal nerve cord), the bottom line (ventral nerve cord), and the wild-type PVD axon and primary dendrite (green) are shown.

We isolate mutants through genetic screens that display profoundly affected dendrite branching and self-avoidance in PVD neurons. We identify mutations in the *kpc-1* gene as being responsible for defective dendrite branching and self-avoidance phenotypes through whole genome sequencing. *kpc-1* mutants display a significantly lower tertiary branch avoidance index and a decreased sensory coverage in the skin compared to wild-type animals. Although *kpc-1* has been implicated in dendrite self-avoidance and arborization, the mechanism by which *kpc-1* regulates these processes is unknown(*4*, *22*, *23*). KPC-1 encodes a proprotein convertase subtilisin/kexin (PCSK). Here, we show that *kpc-1* functions in PVD neurons to regulate dendrite branching and self-avoidance via the *kpc-1* 3’UTR. The *kpc-1* 3’UTR is required for *kpc-1* mRNA localization to distal and higher-order dendrites and promotes its local translation. Although over-expression of *kpc-1* causes greater self-avoidance, it does not limit initial dendrite outgrowth, which supports a direct role for *kpc-1* in self-avoidance. *dma-1* over-expression also displays similar secondary dendrite branching and tertiary dendrite self-avoidance defects that are suppressed with *kpc-1* over-expression. Thus, DMA-1 is a potential KPC-1 target that becomes down-regulated with increased KPC-1 activity. Our results suggest a model in which KPC-1 proteins are synthesized at branching points and contact points between sibling dendrites, leading to local down-regulation of DMA-1 receptors and the consequent dendrite branching and self-avoidance.

## Results

### Isolation of mutants that display profound self-avoidance defects in dendrites of PVD neurons

*C. elegans* PVD neurons extend a large and highly branched dendrite arbor directly beneath the hypodermal surface that envelops the worms to mediate an avoidance response to harsh mechanical force as well as to other noxious stimuli including temperature(*2*). The elaborate dendrite branching patterns that PVD neurons display are constrained by two organizing principles: dendrite branching(*4*) and dendrite self-avoidance. Self-avoidance is an important neuronal process in which dendritic branches arising from a single soma (also called isoneuronal) turn away from one another to minimize crossing and overlap and to promote a uniform receptive field(*24*).

The self-avoidance in *C. elegans* is likely to provide new mechanistic insight into the process as the well-known self-avoidance molecule Dscam(*9–13*)is absent from the *C. elegans* genome. In order to uncover new mechanisms responsible for self-avoidance in *C. elegans*, we screened approximately 9,500 genomes for mutants that displayed self-avoidance defects in secondary and tertiary dendrites. These screens uncovered three strong mutants (*xr47*, *xr58*, and *xr60*) that show profound self-avoidance defects in all animals examined (n>200 each). *xr47* and *xr60* mutants showed an additional secondary dendrite branching phenotype (a defect in the dendrite branching), which resulted in significant reduced number of secondary branches in *xr47* and *xr60* mutants compared to wild-type animals (Table 1). Despite displaying extensive defects in self-avoidance among secondary and tertiary sibling dendrites, *xr58* mutants showed little secondary dendrite branching defect (Fig. 1 a, c; Table 1). In wild-type animals, secondary dendrites that initially contacted neighboring secondary branches retracted in order to form a regularly spaced array of parallel branches (Fig. 1 a, b). In *xr58* mutants, many secondary dendrites remained in contact with each other (Fig. 1c). In wild-type animals, tertiary dendrites initially extended but later retracted once their termini come into contact with sibling tertiary branches (Supplementary Fig. 1). In *xr58* mutants, many tertiary dendrites remained in contact with their siblings long after arborization ended (Fig. 1c; Supplementary Fig. 1), resulting in a significantly lower tertiary branch avoidance index (TBAI) compared to the wild-type animals (Fig. 1d; TBAI: 0.38 in *xr58* mutants versus 0.93 in wild-type, p<0.0001). In wild-type animals, PVD dendrite arbors were fully extended beneath the hypodermis and covered the areas from the posterior animal body to right before the head region (Fig. 1f, Supplementary Fig. 2a, 2c, and 2d). In *xr58* mutants, PVD dendrite arbors developed partially overlapping menorahs, causing decreased sensory coverage of skin (Fig. 1f, Supplementary Fig. 2b, 2c, and 2d). We quantified sensory coverage of skin by PVD dendrite arbors using two measurements, receptive field index and length ratio of receptive field (Supplementary Fig. 2c, d). Both measurements showed significantly reduced values in *kpc-1(xr58)* mutants compared to the wild-type animals (Supplementary Fig. 2).

**Figure 8**
| Genotype | N | Menorah coverage zone<br>per 200 $\mu\text{m}$ of 1° dendrite | Trapped 2° dendrite zone<br>per 200 $\mu\text{m}$ of 1° dendrite |
| --- | --- | --- | --- |
| Wild type | 32 | 183.99 | 0 |
| *kpc-1(xr58)* | 39 | 182.88 | 38.51 |
| *kpc-1(xr47)* | 32 | 143.46 | 73.18 |
| *kpc-1(xr60)* | 27 | 58.73 | 163.90 |
| *kpc-1(gk8)* | 32 | 3.14 | 196.86 |

### Identification of responsible mutations in the *kpc-1* gene

We identified *xr47*, *xr58*, and *xr60* mutations in the *kpc-1* gene through whole genome sequencing. The *xr58* allele harbors a missense mutation resulting in a P440S amino acid substitution at a conserved proline residue four amino acids away from the catalytic serine residue toward the C-terminal(*4*). It was shown by others that *kpc-1(my24)*, a different mutation in the same residue (P440L), caused a loss of function in the *kpc-1* gene which resulted in defective secondary dendrite branching in PVD neurons(*22*). Thus, it appears that mutations at P440 could alter KPC-1 functions moderately or severely. Using the complementation test, we showed that the *kpc-1(gk8)* null mutation failed to complement the *xr58* mutant phenotype in PVD neurons, further confirming that *xr58* is a new *kpc-1* allele (Fig. 6d). The *xr58* mutant reveals an additional function of *kpc-1* in tertiary dendrite self-avoidance as trans-heterozygous mutants (*xr58/gk8*) display the *xr58* mutant phenotype of defective tertiary dendrite self-avoidance rather than defective secondary dendrite branching characteristic of *gk8* (Fig. 1c, 2d). The two other *kpc-1* alleles are also predicted to affect KPC-1 protein structure and function. The *xr60* allele contains a missense mutation, which results in a G223S amino acid substitution in the catalytic domain, whereas the *xr47* allele contains a G-to-A transition at the +1 position of intron 5, which abolishes the donor splice site (Fig. 1e). Compared to the *gk8* null allele, *xr58*, *xr47*, and *xr60* alleles are weaker in their phenotypic effects in PVD neurons as judged by the extent of menorah coverage and trapped secondary dendrites along the primary dendrite axis (Table 1). *kpc-1* mRNAs can be made from *xr58*, *xr47*, and *xr60* mutant alleles based on RT-PCR amplification of respective *kpc-1* cDNAs.

### Elements in the *kpc-1* 3’UTR are required for dendrite self-avoidance

The *kpc-1* genomic DNA containing the 5’, coding, and 3’ fragments rescued both defective secondary dendrite branching and defective tertiary dendrite self-avoidance in the *kpc-1(gk8)* null allele as well as defective tertiary dendrite self-avoidance in the *kpc-1(xr58)* allele (Fig. 2 a, b). To investigate the genomic requirement of the *kpc-1* gene in different steps of PVD dendrite development, we first reduced the *kpc-1* gene to the *kpc-1* coding region without introns. We found that the *kpc-1* gene without introns led to a complete rescue of the dendrite phenotypes in both *kpc-1(gk8)* null and *kpc-1(xr58)* alleles (Fig. 2 a, b). We then further reduced the *kpc-1* promoter to a PVD cell specific promoter (The F49H12.4 promoter). This *Ppvd::kpc-1::kpc-1 3’UTR* transgene remained able to cause a near complete phenotypic rescue, in a manner similar to the *kpc-1* genomic DNA, in both *kpc-1(gk8)* null and *kpc-1(xr58)* alleles, indicating that *kpc-1* acts cell-autonomously in PVD (Fig. 2a, b, and e-h). Consistent with a cell-autonomous role of *kpc-1* in PVD, the expression of the *kpc-1* promoter reporter was detected in PVD neurons during secondary dendrite branching and tertiary dendrite self-avoidance (Fig. 2c). When the *kpc-1* 3’UTR was replaced with an unrelated *unc-54* 3’UTR, the *Ppvd::kpc-1::unc-54 3’UTR* transgene was able to largely, albeit incompletely, rescue defective secondary dendrite branching in the *kpc-1(gk8)* null allele (Fig. 2 a, b, d). It failed to rescue defective tertiary dendrite self-avoidance in the *kpc-1(gk8)* null allele (Fig. 2 a, b, d), suggesting a more stringent 3’UTR requirement for tertiary dendrite self-avoidance. In addition to tertiary dendrites, some secondary dendrites displayed self-avoidance defects in these transgenic animals (Fig. 2d). The same *Ppvd::kpc-1::unc-54 3’UTR* transgene was also unable to rescue the defective tertiary dendrite self-avoidance in the *kpc-1(xr58)* allele (Fig. 2b). Increasing the concentration of the *Ppvd::kpc-1::unc-54 3’UTR* transgene from 10 ng/μl to 100 ng/μl did not alter this outcome. Thus, these results show that tertiary dendrite self-avoidance depends on elements in the *kpc-1* 3’UTR.

**Figure 2.**
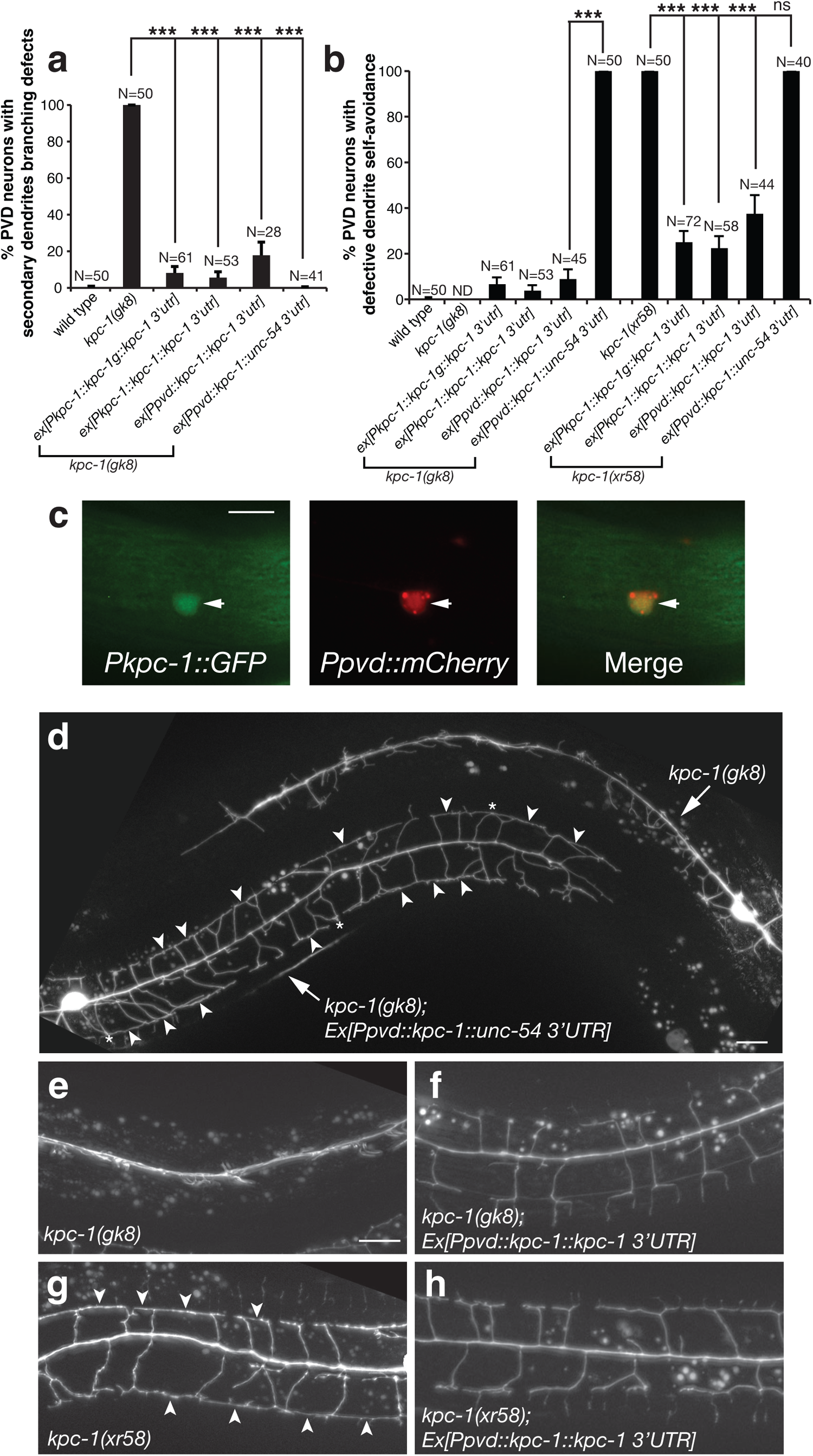
*kpc-1* is required for both secondary dendrite branching and tertiary dendrite self-avoidance but the requirements of two functions are different. (a) *kpc-1* functions in PVD neurons to promote secondary dendrite branching largely independent of the *kpc-1* 3’UTR. Percentage PVD neurons with secondary dendrite branching in *kpc-1(gk8)* mutants carrying various transgenes. ***p < 0.001 by a two-proportion *Z*-test. (b) The *kpc-1*’s function in PVD for tertiary dendrite self-avoidance is dependent on the *kpc-1* 3’UTR. Percentage PVD neurons with defective tertiary dendrite self-avoidance in *kpc-1(gk8)* and *kpc-1(xr58)* mutants carrying various transgenes. ND, not determined. ***p < 0.001 by a two-proportion *Z*-test. (c) Identification of the *kpc-1* promoter reporter expression in PVD neurons. (d) Images of PVD neurons in *kpc-1(gk8)* mutants with or without the *Ppvd::kpc-1::unc-54 3’UTR* transgene. The *Ppvd::kpc-1::unc-54 3’UTR* transgene rescued the secondary dendrite branching but not the tertiary dendrite self-avoidance defect in *kpc-1(gk8)* mutants. Arrowheads point to contacts between neighboring tertiary dendrites. Asterisks indicate contacts between neighboring secondary dendrites. (e, f) Images of PVD neurons in *kpc-1(gk8)* mutants without or with the *Ppvd::kpc-1::kpc-1 3’UTR* transgene. The *Ppvd::kpc-1::kpc-1 3’UTR* transgene rescued both the secondary dendrite branching and the tertiary dendrite self-avoidance defect in *kpc-1(gk8)* mutants. (g, h) Images of PVD neurons in *kpc-1(xr58)* mutants without or with the *Ppvd::kpc-1::kpc-1 3’UTR* transgene. The *Ppvd::kpc-1::kpc-1 3’UTR* transgene rescued the tertiary dendrite self-avoidance defect in *kpc-1(xr58)* mutants. Arrowheads point to contacts between neighboring tertiary dendrites. Scale bar, 20 μm.

### The *kpc-1* 3’UTR is required for localization of *kpc-1* mRNAs to distal primary dendrites and higher-order tertiary dendrites in PVD neurons

To directly visualize *kpc-1* mRNA distribution in PVD neurons, we utilized the MS2 tagging system, which relies on the sequence-specific interaction between the MS2 bacteriophage RNA hairpin loops and capsid proteins. In this system, the *kpc-1* mRNA containing either its own 3’UTR or a control *unc-54* 3’UTR was fused to tandem MS2 binding sites and co-expressed with a green fluorescent MS2 capsid fusion protein(*25*)(*26*) (Fig. 3a). The green fluorescent MS2 capsid fusion protein, which is targeted to the nucleus by the nuclear localization signal (Fig. 3b), binds tightly to MS2 binding sites fused to the 3’UTR of the *kpc-1* mRNA, allowing visualization of the *kpc-1* mRNA by MS2 fluorescence in PVD neurons. Using this system, we detected significant axonal and dendritic signals for *kpc-1* transcripts containing its own 3’UTR (Fig. 3c, d, g). Dendritic signals were preferentially distributed in the 1° and 3° dendrites (Fig. 3c, d), consistent with *kpc-1*’s roles in regulating secondary dendrite branching from the 1° dendrites and tertiary dendrite self-avoidance. Distribution of *kpc-1* transcripts in 3° dendrites was region specific. We observed enriched localization at potential branching points and retracted contact points between neighboring 3° dendrites (Fig. 3c, d). Using the mCherry reporter to label PVD dendrites, we can confirm enrichment of *kpc-1* transcripts at branching points and retracted contact points (Fig. 3e). In contrast, when the *kpc-1* 3’UTR was replaced with an unrelated *unc-54* 3’UTR, the *kpc-1* transcripts were mainly trapped in the nucleus with some perinuclear signals detected in PVD (Fig. 3f). *kpc-1* transcripts containing the *unc-54* 3’UTR had a significantly lower chance to be transported to higher-order tertiary dendrites compared to *kpc-1* transcripts containing its own 3’UTR (Fig. 3g, h). Together, these results indicate that the *kpc-1* 3’UTR facilitates *kpc-1* RNA localization to distal primary dendrites and higher-order tertiary dendrites, implicating local translation of *kpc-1* in dendrites.

**Figure 3.**
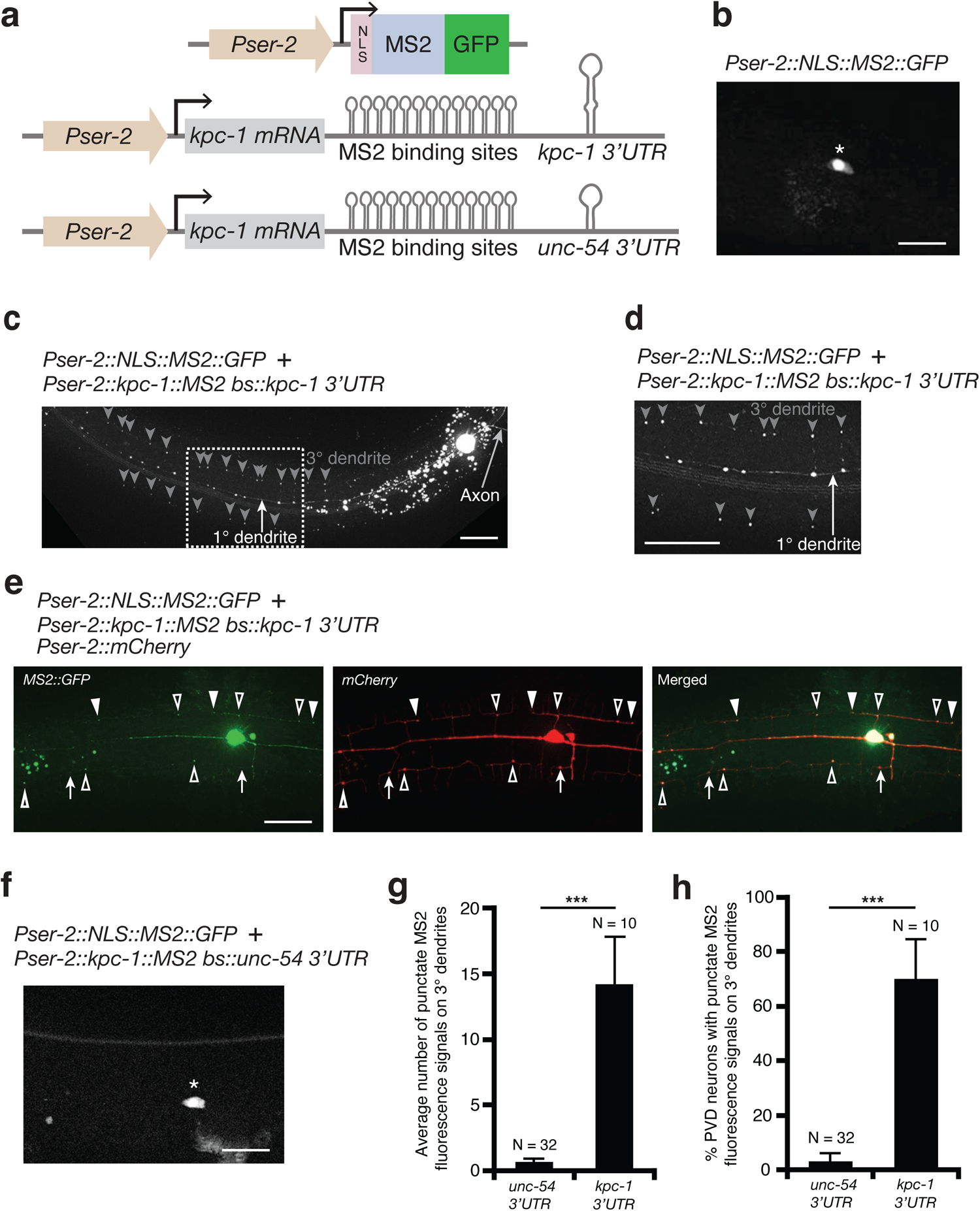
The *kpc-1* 3’UTR targets the *kpc-1* mRNA to higher-order dendrites. (**a**) The MS2 tagging system is composed of the *kpc-1* mRNA containing its own 3’UTR or a control *unc-54* 3’UTR fused to 6xMS2 binding sites co-expressed with a green fluorescent MS2 capsid protein. The nuclear localization signal (NLS) is used to initially target green fluorescent MS2 proteins to the nucleus. (**b**) Image of the PVD neuron showing enriched nuclear signals for the nuclear localized green fluorescent MS2 capsid protein. Asterisk indicates soma. Scale bar, 20 μm. (**c**) Image of the PVD neuron showing axonal and dendritic signals for *kpc-1* transcripts containing the *kpc-1* 3’UTR and MS2 binding sites. The white arrow points to dendritic signals in the primary dendrite and the grey arrow marks axonal signals. Grey arrowheads indicate the potential branching points at 3° dendrites and contact points between neighboring 3° dendrites. The dashed box area in 3**c** was blown up and shown in 3**d**. Anterior is to the left, dorsal is up. Scale bar, 20 μm. (**e**) Image of the PVD neuron showing axonal and dendritic signals for *kpc-1* transcripts containing the *kpc-1* 3’UTR and MS2 binding sites. Low level of the mCherry protein was used to label PVD. Arrowheads point to potential tertiary dendrite contact points and open arrowheads denote dendrite branching points. Arrows show red dendrite markers that are not overlapped with any green MS2::GFP signals, indicating that red signals did not bleed through to give rise to green signals. Scale bar, 20 μm. (**f**) Image of the PVD neuron showing enriched soma signals for *kpc-1* transcripts containing the control *unc-54* 3’UTR and MS2 binding sites. Asterisk indicates soma. Scale bar, 20 μm. (**g**) Average number of punctate MS2 fluorescence signals on 3° dendrites in transgenic animals expressing *kpc-1* transcripts, containing either its own 3’UTR or the control *unc-54* 3’UTR. ***p < 0.001 by a Student’s *t*-test. (**h**) Percentage PVD neurons with punctate MS2 fluorescence signals on 3° dendrites in transgenic animals expressing *kpc-1* transcripts, containing either its own 3’UTR or the control *unc-54* 3’UTR. ***p < 0.001 by a two-proportion *Z*-test.

### The *kpc-1* 3’UTR promotes local protein synthesis in the distal segment of PVD dendrites

Our results showed that the *kpc-1* 3’UTR facilitates localization of *kpc-1* transcripts to the distal primary and tertiary PVD dendrites, suggesting that the *kpc-1* 3’UTR promotes local translation of transcripts it regulates. To examine whether the *kpc-1* 3’UTR can promote local protein synthesis, we performed fluorescence recovery after photoconversion (FRAP) of the photoconvertible Kaede protein(*27*), which was under control of the *kpc-1* 3’UTR or a control *unc-54* 3’UTR. Upon photostimulation using 405 nm laser, Kaede proteins can be converted from green fluorescent proteins to red fluorescent proteins irreversibly, allowing us to distinguish the future newly synthesized green fluorescent Kaede proteins from the resident (converted) red fluorescent Kaede proteins. We carried-out photoconversion at a broad region from the mid-body to the nerve ring and measured newly synthesized green fluorescent Kaede proteins in a local area distant away from the cell body (Fig. 4a). This design ensures that the local green fluorescence signal represents newly synthesized Kaede proteins rather than resident (unconverted) Kaede proteins diffused from the soma (Fig. 4a). Green fluorescence recovery of Kaede proteins expressed from the *kaede::kpc-1* 3’UTR is significantly faster than that from the *kaede::unc-54* 3’UTR (Fig. 4b, c), indicating that the *kpc-1* 3’UTR promotes local protein synthesis in the distal segment of PVD dendrites. Using the split GFP *rps-18* reporter to visualize ribosomes in PVD neurons(*28*), we detected the protein synthesis machinery not only in the soma but also in secondary and tertiary dendrites (Fig. 4d), further supporting protein synthesis occurs in the higher order dendrites to allow for proteome remodeling.

**Figure 4.**
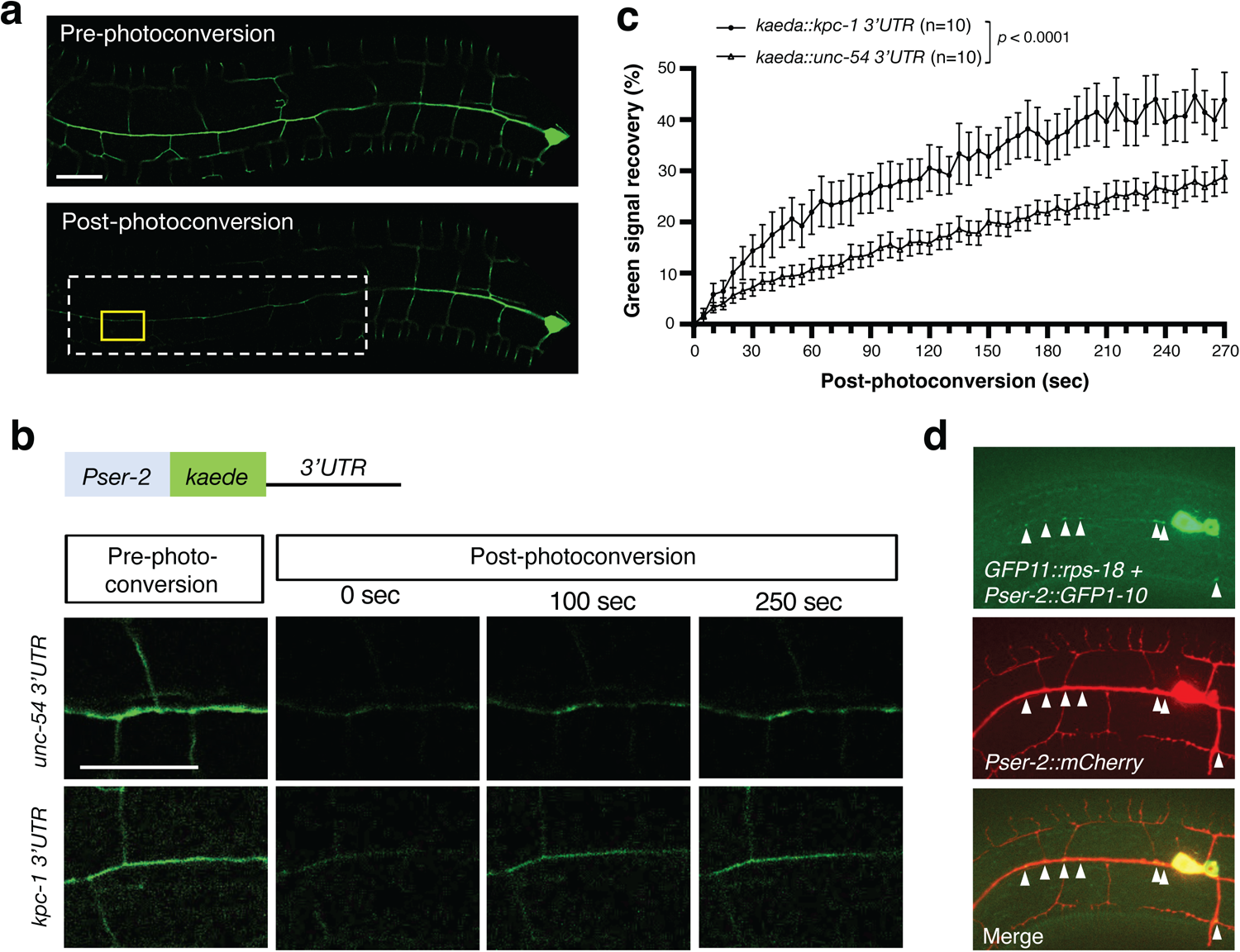
The *kpc-1* 3’UTR promotes local protein synthesis in the distal segment of PVD dendrites. (**a**) Representative images of Kaeda-labelled PVD dendrites pre- and post-photoconversion. The dashed white box indicates a wide photoconverted region. The solid yellow box denotes a targeted area of analysis, which is distal to the soma within the wide photoconverted region. Scale bar, 20 μm. (**b**) Pre-photoconversion of resident Kaede proteins and post-photoconversion of newly synthesized Kaede proteins. Kaeda expression in PVD dendrites is controlled by either a control *unc-54* 3’UTR or a *kpc-1* 3’UTR. Scale bar, 20 μm. (**c**) The *kaeda::kpc-1 3’UTR* transcript is expressed at a higher rate compared to the control *kaeda::unc-54 3’UTR* transcript. Quantification of green signal recovery in a targeted distal area after photoconversion of a wider region of PVD dendrites. Statistical analysis using 2-way ANOVA. Average data of Kaeda expression intensity are presented as means ± SEM. (**d**) Expression of a split GFP reporter allows visualization of ribosomes in PVD neurons. The smaller part of split GFP (GFP11) is tagged to the endogenous *rps-18* (GFP11::rps-18). The larger part of split GFP (GFP1-10) is overexpressed from a PVD promoter *ser-2* using a high-copy transgene. GFP1-10 binds to the GFP11::RPS-18 in neurons and illuminates RPS-18. The *Pser-2::mCherry* marker labels PVD dendrites. Ribosomes are mostly restricted in the PVD soma, which is marked by an asterisk. Arrowheads indicate RPS-18 signals. Scale bar, 20 μm.

### Identification and characterization of secondary structural motifs in the *kpc-1* 3’UTR required for tertiary dendrite self-avoidance

Many studies have shown that specific elements in 5′ or 3′ untranslated regions (UTRs) of mRNA regulate mRNA stability, splicing, translation and localization(*29*). microRNAs regulate mRNA translation and stability by binding to complementary sequences in the 3′UTR(*30*). Other types of elements in the 3’UTR can function in either a sequence- or secondary structure-specific manner. For example, one of the earliest identified localization elements was discovered from the chick β-actin mRNA. The “zipcode” in the 3′UTR of β-actin mRNA consists of 54 nucleotides that contain tandem repeats of the conserved hexanucleotide motif ACACCC and form a stem-loop structure recognized by the Zipcode Binding Proteins (ZBPs) to regulate the localization and translation of β-actin mRNA(*31*, *32*). In yeast, the cis-acting sequences that mediate ASH1 mRNA targeting to the bud tip provide another example of repetitive clustering and synergistic action of localization elements. Four localization elements in ASH1 mRNA were all predicted to form stem-loop structures, and each element on its own is able to localize mRNAs, but the presence of four elements together increases the efficiency of localization(*33–35*). In Aplysia sensory-motor neurons, cis-elements in the 3′ UTR of the neurotransmitter sensorin mRNA control the transport of sensorin mRNAs from soma to neurite(*36*). In mice, conditional deletion of an axon-localizing cis-element in the 3′UTR of importin β1 mRNAs in sensory neurons caused selective depletion of importin β1 mRNAs and proteins in axons, without affecting cell body levels or nuclear functions, suggesting that local translation of importin β1 mRNAs in axons enables separation of axonal and nuclear transport functions of importins, and is required for efficient retrograde signaling in injured axon(*37*).

In *C. elegans*, it was previously reported that the *cebp-1* 3’UTR is required for the localization of *cebp-1* mRNAs to discrete locations in touch axons, which can be translated into proteins locally(*38*). What is unclear is whether those discrete locations represent synapses in touch axons. In addition, a putative stem-loop secondary structure was identified in the *cebp-1* 3’UTR that inhibits *cebp-1* mRNA stability or its translation(*39*). Thus, the *cebp-1* 3’UTR appears to have both positive and negative impacts on *cebp-1* gene expression.

Secondary structural motifs derived from the alignment of *kpc-1* 3’UTRs from 4 closely-related nematode species, including *C. elegans* (Supplementary Fig. 3), as well as individual Turbofold II-predicted secondary structures(*40*), revealed that two stem-loop structural motifs (SLS1 and SLS2) are conserved in the 3’ UTR of all *kpc-1* homologs (Fig. 5a). The *kpc-1* transgene with its 3’UTR being replaced by a control *unc-54* 3’UTR lost the ability to rescue tertiary dendrite self-avoidance defects in the *kpc-1(gk8)* null allele (Fig. 5f). Deletion analysis showed that SLS2 but not SLS1 is required for the *kpc-1* 3’UTR to mediate tertiary dendrite self-avoidance since *ΔSLS2* but not *ΔSLS1* in the *kpc-1* 3’UTR abolished the ability of the *kpc-1* transgene to rescue tertiary dendrite self-avoidance defects in the *kpc-1(gk8)* null allele (Fig. 5f). The *kpc-1* transgene with mutations that disrupt base-pairing in the stem region of SLS2 in the 3’UTR exhibited significantly more contacts between sibling tertiary dendrites (indicating losing self-avoidance functions; Fig. 5c, f) whereas the *kpc-1* transgene with reciprocal mutations that preserve base-pairing in the stem region of SLS2 in the 3’UTR did not cause more contacts between sibling tertiary dendrites (indicating maintaining self-avoidance functions; Fig. 5d, f).

**Figure 5.**
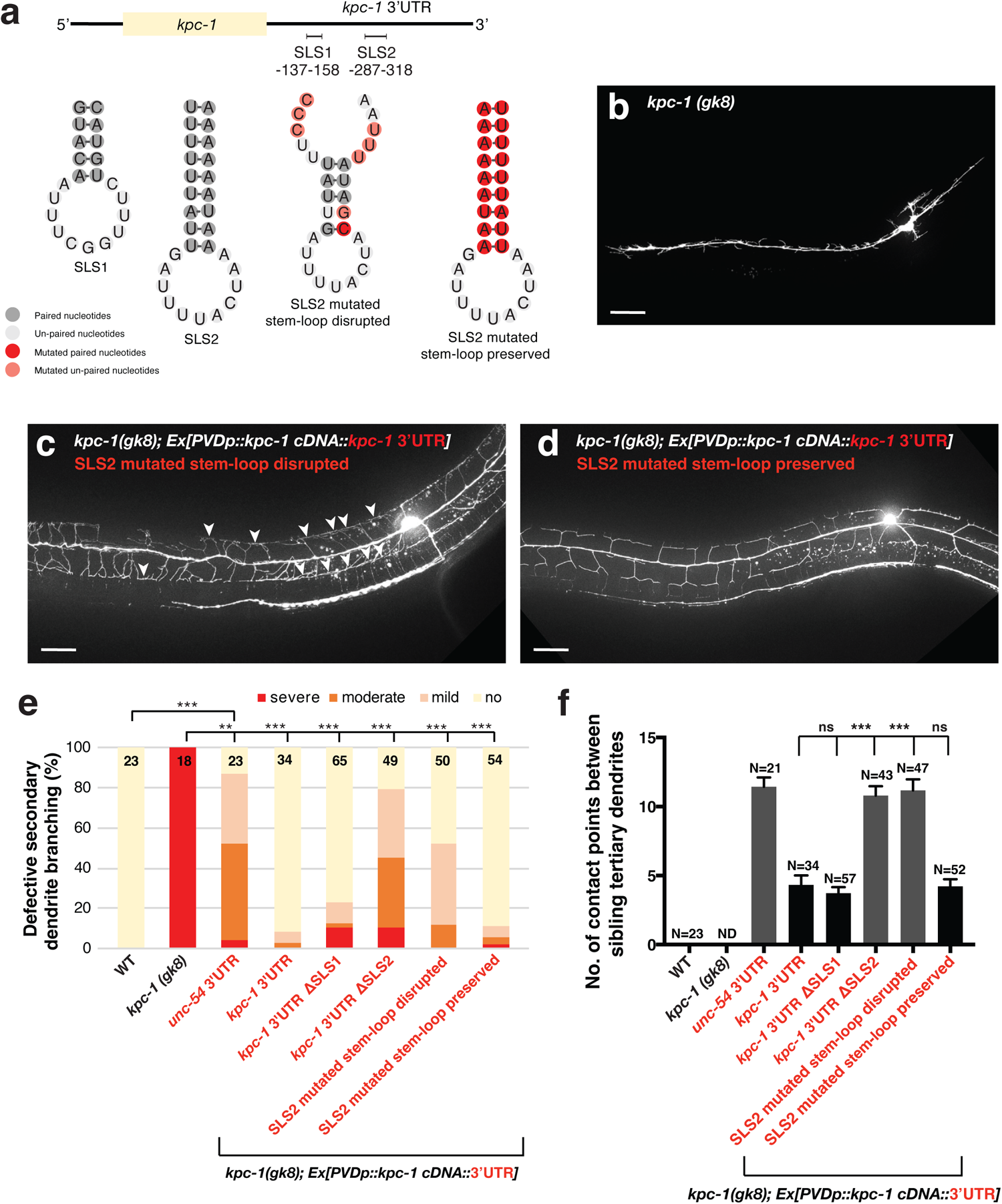
A putative stem-loop secondary structure in the *kpc-1* 3’UTR is required for tertiary dendrite self-avoidance. (**a**) Illustration of putative stem-loop secondary structural motifs and specific mutations in the *kpc-1* 3’UTR. (**b-d**) Representative images of the *kpc-1(gk8)* mutant, the *kpc-1(gk8)* mutant expressing the *Ppvd::kpc-1::kpc-1* 3’UTR transgene (SLS2 mutated base pairing disrupted), and the *kpc-1(gk8)* mutant expressing the *Ppvd::kpc-1::kpc-1* 3’UTR transgene (SLS2 mutated base pairing preserved). Arrowheads point to contacts between neighboring tertiary dendrites. Scale bars, 20 μm. (**e**) Percentage PVD neurons with defective secondary dendrite branching in wild-type, *kpc-1(gk8)* mutants, and *kpc-1(gk8)* mutants carrying various transgenes. **p < 0.01 and ***p < 0.001 by a two-proportion *Z*-test. (**f**) Quantification of contact points between sibling tertiary dendrites in wild-type and *kpc-1(gk8)* mutants carrying various transgenes. ND, not determined. \*\*\**p*<0.001 by one-way ANOVA with Dunnett’s test.

All the *kpc-1* transgenes tested here partially rescued secondary dendrite branching defects in the *kpc-1(gk8)* null allele, suggesting that the modifications we introduced to the transgenes, including the 3’UTR replacement, deletions, and mutations, did not result in degradation of the *kpc-1* transcripts (Fig. 5e). Levels of the *kpc-1* RNA expressed from various transgenes in *kpc-1(gk8)* mutants were analyzed by RT-PCR. All *kpc-1* 3’UTR modifications were made from the PF49H12.4::*kpc-1s*::*kpc-1* 3’UTR construct, where only the short-isoform *kpc-1s* RNA would be generated. We found that the *kpc-1* RNA expression levels are comparable among different *kpc-1* transgenes (Supplementary Fig. 4), indicating that modifications we introduced to the *kpc-1* 3’UTR did not affect the *kpc-1* RNA stability. Furthermore, even though the level of the endogenous *kpc-1* RNA in wild-type animals (containing both long and short isoforms) is lower, it promotes dendrite self-avoidance better than a higher level of the short-isoform (*kpc-1s* RNA) expressed from various transgenes in PVD neurons of *kpc-1(gk8)* mutants (Supplementary Fig. 4). Together, our results have identified an important secondary structural motif in the *kpc-1* 3’UTR required for tertiary dendrite self-avoidance.

To validate the role of the *kpc-1* 3’UTR secondary structural motif SLS2 in PVD dendrite branching and self-avoidance, we applied the CRISPR-Cas9 engineering to generate mutations in the *kpc-1* 3’UTR. Due to the AT-rich sequence in the region surrounding the SLS2 site, only suboptimal sgRNAs with low target selectivity could be designed, which prevented generation of specific SLS2 mutations. A more distal but selective sgRNA was subsequently designed to generate a 296 base-pair deletion that includes the SLS2 site. Although transgene experiment using *kpc-1(gk8)* mutants carrying the *Ppvd::kpc-1::kpc-1* 3’UTR transgene with mutated SLS2 base pairing demonstrated an importance role of SLS2 in dendrite self-avoidance, the CRISPR-Cas9 engineered *kpc-1(xr82) [SLS2 deleted]* mutant allele only showed a weak tertiary dendrite self-avoidance defect (Supplementary Fig. 5a). The mild phenotypic effects of the *kpc-1(xr82)* mutant allele were inadvertently caused by the creation of a new stem-loop structure that likely substitutes the SLS2’s function. Although the *xr58* or *xr82 [SLS2 deleted]* allele alone did not cause a severe self-avoidance defect, introducing the *xr84 [SLS2 deleted]* allele into the *kpc-1(xr58)* background collectively caused significantly more secondary dendrite branching defects compared to wild-type (Supplementary Fig. 5b). Taken together, our results indicate that both the *kpc-1* coding region and its 3’UTR are required for PVD dendrite patterning.

### KPC-1 down-regulates DMA-1 receptors on PVD tertiary dendrites

While the number of tertiary dendrites was significantly reduced in *sax-7*, *mnr-1*, *lect-2*, or *dma-1* mutants, it was significantly increased in *kpc-1(xr58)* mutants(*16*, *17*, *19*– *21*) (Fig. 1c). These results suggested that *kpc-1* likely antagonizes the *sax-7/mnr-1/lect-2/dma-1* signaling pathway. Double mutant analyses revealed that *sax-7*, *mnr-1*, *lect-2*, and *dma-1* mutations completely suppressed the *kpc-1(xr58)* mutant phenotype of excessive tertiary dendrites, indicating that the *kpc-1* mutant effect depends on the *sax-7/mnr-1/lect-2/dma-1* signaling pathway. The *kpc-1(gk8)* null allele seems to have a phenotype very similar to the *sax-7*, *mnr-1*, *lect-2*, and *dma-1* mutants(*16*, *17*, *19–21*) (Fig. 2d, e). However, closer examination of the *kpc-1(gk8)* null mutant phenotype revealed that defective secondary dendrite extension to sublateral longitudinal stripes is caused by a failure of secondary dendrites branching out(*4*), a phenotype that is clearly different from the less secondary dendrite outgrowth in *sax-7*, *mnr-1*, *lect-2*, and *dma-1* mutants. Double mutant analyses between the *kpc-1(gk8)* null allele and *sax-7*, *mnr-1*, and *dma-1* mutations have been reported in a recent paper(*4*), which also supported that *kpc-1* antagonizes the *sax-7*/*mnr-1*/*dma-1* signaling pathway in PVD neurons. *dma-1*, like *kpc-1*, encodes a transmembrane protein and acts cell-autonomously in PVD neurons(*4*, *19*). Thus, DMA-1 could be a potential KPC-1 target in regulating PVD tertiary dendrite self-avoidance. Supporting this possibility, endogenous DMA-1 protein levels on PVD tertiary dendrites were up-regulated in *kpc-1* mutants, suggesting that KPC-1 normally down-regulates DMA-1 on PVD tertiary dendrites (Fig. 6 a, b). In addition, the *kpc-1(xr47)* mutant allele, which showed a greater increase in the number of overlapped tertiary dendrites compared to the *kpc-1(xr58)* mutant allele (Fig. 6c), caused a greater up-regulation of the DMA-1 protein level on PVD tertiary dendrites than the *kpc-1(xr58)* mutant allele (Fig. 6 a, b). A recent paper reported a different phenotypic effect of *kpc-1* mutations in PVD neurons where *kpc-1* mutations caused excessive DMA-1 receptors on secondary dendrites, which resulted in secondary dendrites trapped in the primary dendrite zone where a high level of the cognate ligand SAX-7/L1CAM is present(*4*).

**Figure 6.**
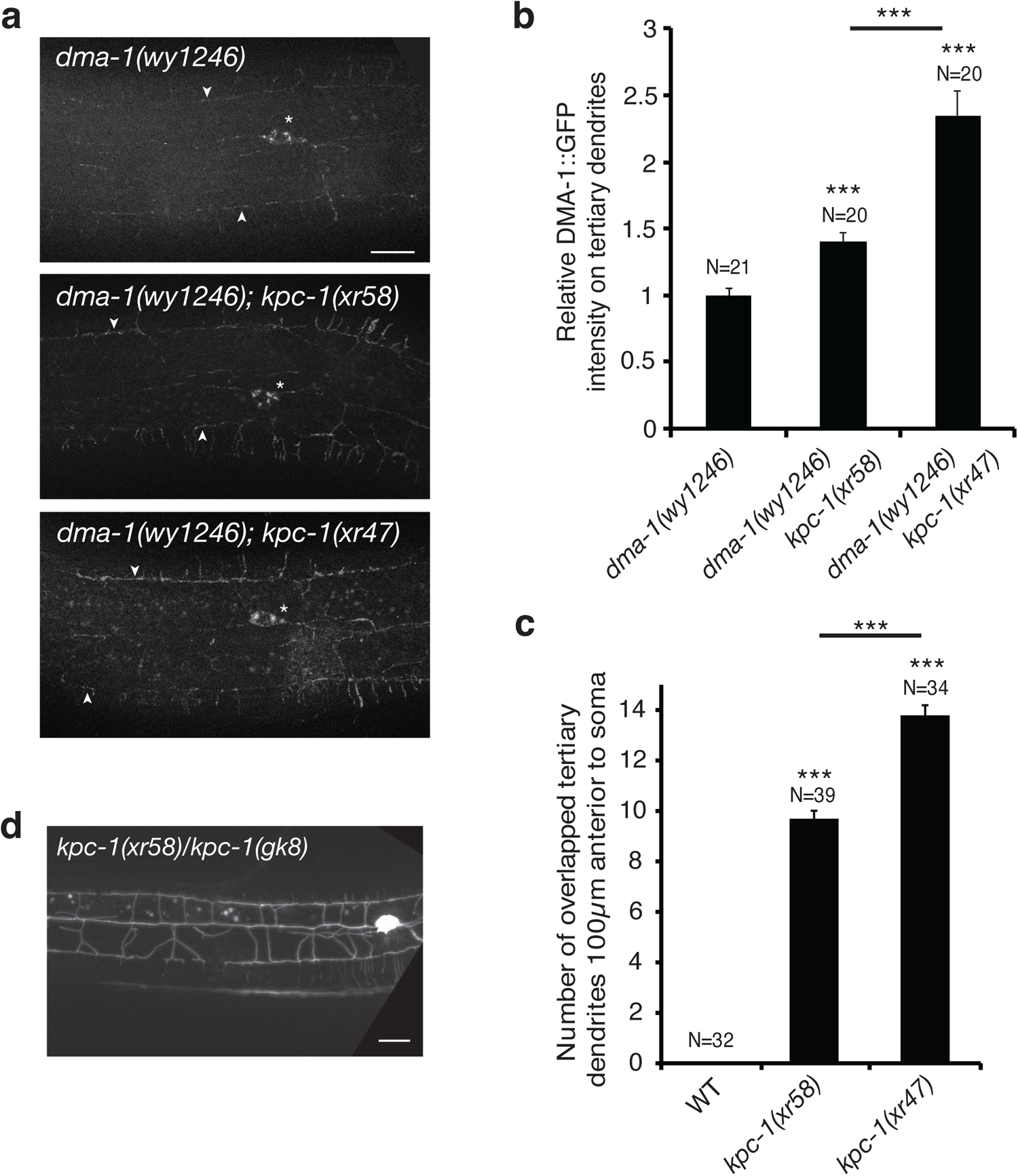
*kpc-1* alleles differentially affect DMA-1 levels on 3° dendrites and 3° dendrite self-avoidance. (**a**) Representative images showing DMA-1::GFP expression on PVD dendrites in wild type, *kpc-1(xr58)*, and *kpc-1(xr47)* mutants. Asterisk indicates soma. Arrowheads indicate tertiary branches. Scale bars, 20 μm. (**b**) Average fluorescence intensity of the DMA-1::GFP on PVD tertiary dendrites in wild type [*dma-1(wy1246)*], *kpc-1(xr58)* mutants [*kpc-1(xr58); dma-1(wy1246)*], and *kpc-1(xr47)* mutants [*kpc-1(xr47); dma-1(wy1246)*]. The *dma-1(wy1246) [dma-1::GFP]* allele was created by inserting a GFP reporter gene immediately after the transmembrane domain of the endogenous *dma-*1 gene. This reporter strain does not cause any defects in PVD dendrites(*3*). **p < 0.01 and ***p < 0.001 by one-way ANOVA with Tukey’s test. (**c**) Quantification of overlapped tertiary dendrites in wild type versus *kpc-1* mutants. ***p < 0.001 by one-way ANOVA with Tukey’s test. (**d**) Image of the trans-heterozygous *kpc-1(xr58)/kpc-1(gk8)* mutant displaying self-avoidance defects similar to those displayed by homozygous *kpc-1(xr58)* mutants. Scale bars, 20 μm.

### The *kpc-1-dma-1* regulatory circuit promotes dendrite self-avoidance

To further strengthen the conclusion of a *kpc-1-dma-1* regulatory circuit in promoting PVD dendrite self-avoidance, we tested whether *dma-1* over-expression (OE) mimics the *kpc-1* mutation in its phenotypic effect on PVD dendrites. *dma-1* OE displayed a tertiary dendrite self-avoidance defect similar to that exhibited by the *kpc-1(xr58)* mutant allele (Fig. 7 a, b). In addition, *dma-1* OE caused a secondary dendrite branching phenotype (Fig. 7d), which was further enhanced by the *kpc-1(xr58)* mutation. In contrast, *kpc-1* OE suppressed the secondary dendrite branching phenotype in *dma-1* OE animals (Fig. 7c-f). While fewer *kpc-1* OE*; dma-1* OE animals displayed a secondary dendrite branching phenotype, most still exhibited a weak tertiary dendrite self-avoidance defect, which was milder than *dma-1* OE or *kpc-1(xr58)* alone. Thus, a weaker *kpc-1-dma-1* regulatory circuit causes tertiary dendrite self-avoidance defects whereas a further reduced *kpc-1-dma-1* regulatory circuit results in secondary dendrite branching defects. Indeed, the mild *kpc-1(xr58)* mutant allele preferentially affects tertiary dendrite self-avoidance rather than secondary dendrite branching. Together, our results support a model in which locally expressed KPC-1 down-regulates DMA-1 receptors on PVD dendrites to promote dendrite branching and self-avoidance (Fig. 7k).

**Figure 7.**
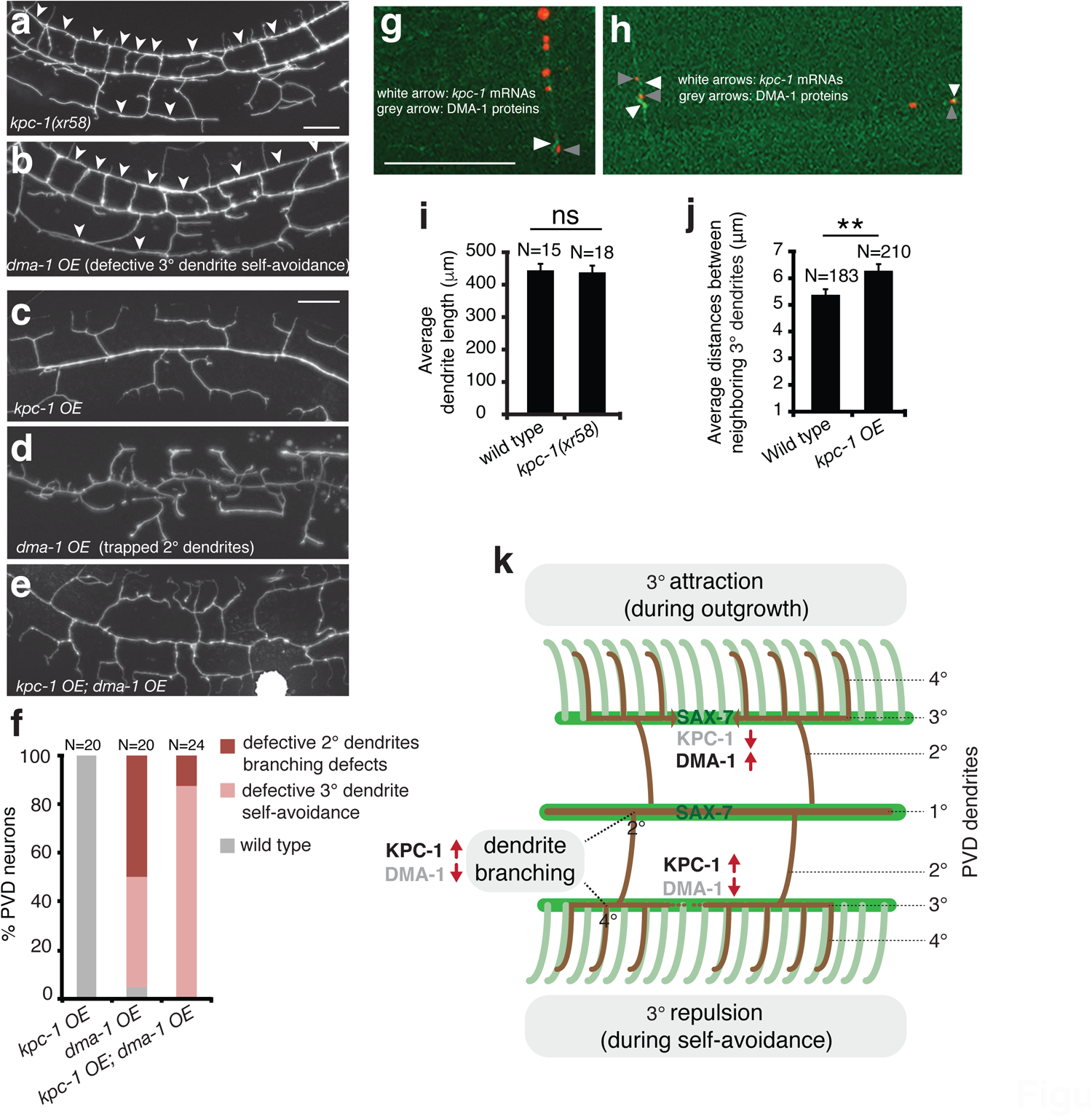
*dma-1* over-expression phenocopied the *kpc-1(xr58)* mutant allele. (a, b) Images of PVD dendrites in *kpc-1(xr58)* mutants and *dma-1* over-expression animals. Arrowheads point to contacts between neighboring tertiary dendrites. Scale bars, 20 μm. (c-e) Images of PVD dendrites in *kpc-1* over-expression animals, *dma-1* over-expression animals, and animals over-expressing both *kpc-1* and *dma-1*. Scale bars, 20 μm. (f) *kpc-1* over-expression suppressed secondary dendrite branching phenotypes in *dma-1* over-expression animals. (g, h) Images of PVD neurons showing dendritic signals for *kpc-1* transcripts and DMA-1 proteins. *kpc-1* transcripts containing *kpc-1* 3’UTR and MS2 binding sites were visualized by green fluorescent MS2 capsid proteins. DMA-1 proteins were visualized by tagged mCherry fluorescent proteins. White arrowheads point to dendritic signals for *kpc-1* transcripts and grey arrowheads indicate dendritic signals for DMA-1 proteins. Scale bars, 15 μm. (i) Bar chart of average PVD dendrite length in wild type and *kpc-1(xr58)* mutants. Dendritomy was carried out at the early L4 stage when tertiary dendrites underwent self-avoidance. Newly grown PVD dendrite length was measured 24 hours after dendritomy. P=0.83 by a Student’s *t*-test. Error bars indicate SEM. (j) Bar chart of average distances between neighboring tertiary dendrites in wild type and *kpc-1* over-expression animals at the young adult stage. For each animal, average distances of the gaps between neighboring tertiary dendrites were measured per 150 μm anterior to the cell body. *kpc-1* over-expression animals differ from wild-type at **p<0.01 by a Student’s *t*-test. Error bars indicate SEM. (k) Model of the *kpc-1-dma-1* regulatory circuit in dendrite branching and self-avoidance. *kpc-1* is expressed at branching points and contact points between the neighboring 3° dendrites, which down-regulates DMA-1 receptors locally, leading to dendrite branching and retraction of sibling dendrites. SAX-7 cues are distributed highly in the 1° and the 3° dendrite growth pathways.

We have made different versions of KPC-1::dzGFP^myr^ and KPC-1::GFP fusion proteins, both wild-type and mutant forms that disrupted the KPC-1 self-cleavage site. However, we were unable to detect any fluorescence signal beyond the PVD cell body (unpublished), suggesting that KPC-1 proteins have very short half-life in PVD dendrites. Due to this technical limitation, we cannot determine whether KPC-1 proteins are synthesized locally in tertiary dendrites. Nor can we address whether the KPC-1 protein and its substrate DMA-1 specifically colocate to higher order dendrites. As an alternative approach, we superimposed the image of dendritic signals for *kpc-1* transcripts with the image of dendritic signals for DMA-1 proteins. We found that they did not show overlapping distribution pattern (Fig. 7 g, h). Instead, adjacent sites of expression can be detected for *kpc-1* mRNAs and DMA-1 proteins (Fig. 7 g, h), consistent with down-regulation of DMA-1 by KPC-1 at specific sites.

The tertiary dendrite self-avoidance defect could result from either excessive dendrite growth ability that overcomes a self-avoidance mechanism or insufficient self-avoidance. To distinguish between these two possibilities, we performed laser dendritomy in PVD neurons at the early L4 stage when tertiary dendrites underwent self-avoidance and assessed the dendrite growth ability 24h after surgery. Dendrites were cut at the primary branch 20 μm anterior to the cell body such that the entire anterior dendrite arbor was disconnected from the cell body after surgery. We found that PVD dendrite growth ability was not significantly enhanced in *kpc-1(xr58)* mutants compared to wild-type animals (Fig. 7i). This result indicated that the dendrite self-avoidance defect in *kpc-1(xr58)* mutants cannot be attributed to excessive dendrite growth ability.

In contrast, over-expression of *kpc-1* significantly increased the distance (gap) between neighboring tertiary dendrites (Fig. 7 c, j). This result suggested that *kpc-1* either limited tertiary branch outgrowth or promoted tertiary dendrite self-avoidance. Time-lapse imaging revealed that neighboring tertiary dendrites in *kpc-1* over-expression animals actually contacted at first and retracted subsequently, suggesting that *kpc-1* over-expression did not limit tertiary branch outgrowth. Together, these results support that *kpc-1* promotes dendrite self-avoidance rather than limiting dendrite outgrowth.

### Defective PVD dendrite arborization causes reduced male mating efficiency

The mechanical sensation is known to regulate animal locomotion and posture during male mating behavior(*41*),(*42*). PVD neurons are known mechanosensors but whether they are involved in male mating behavior is not known. Mature PVD dendrite arbors display a sensory network that innervates the skin area outside of the head region. We characterized PVD dendrites in *kpc-1(xr58)* mutant males and found that *kpc-1(xr58)* mutant males also displayed dendrite self-avoidance defects and reduced sensory coverage of skin by PVD dendrite arbors (Fig. 8a-c). To determine whether fully extended dendrite arbors of PVD neurons are crucial for male mating behavior, we assessed the mating efficiency of *kpc-1* mutant males. The *kpc-1(xr58)* mutants males showed reduced reproductive efficiencies (Fig. 8d), suggesting that PVD neurons are involved in the male courtship process. The male mating process begins with males responding to the hermaphrodite contact, turning around the hermaphrodite’s head and tail, until the vulva is located. We found that *kpc-1(xr58)* mutant males have no problem responding to hermaphrodite contact (Fig. 8e) but are required to make more turns to mate with hermaphrodite successfully (Fig. 8f). Therefore, PVD neurons likely coordinate diverse behavioral motifs in reproductive behavior. These results suggest that male mating behavior requires the mechanosensory inputs from PVD neurons to ensure sufficient mating motor patterns.

**Figure 8.**
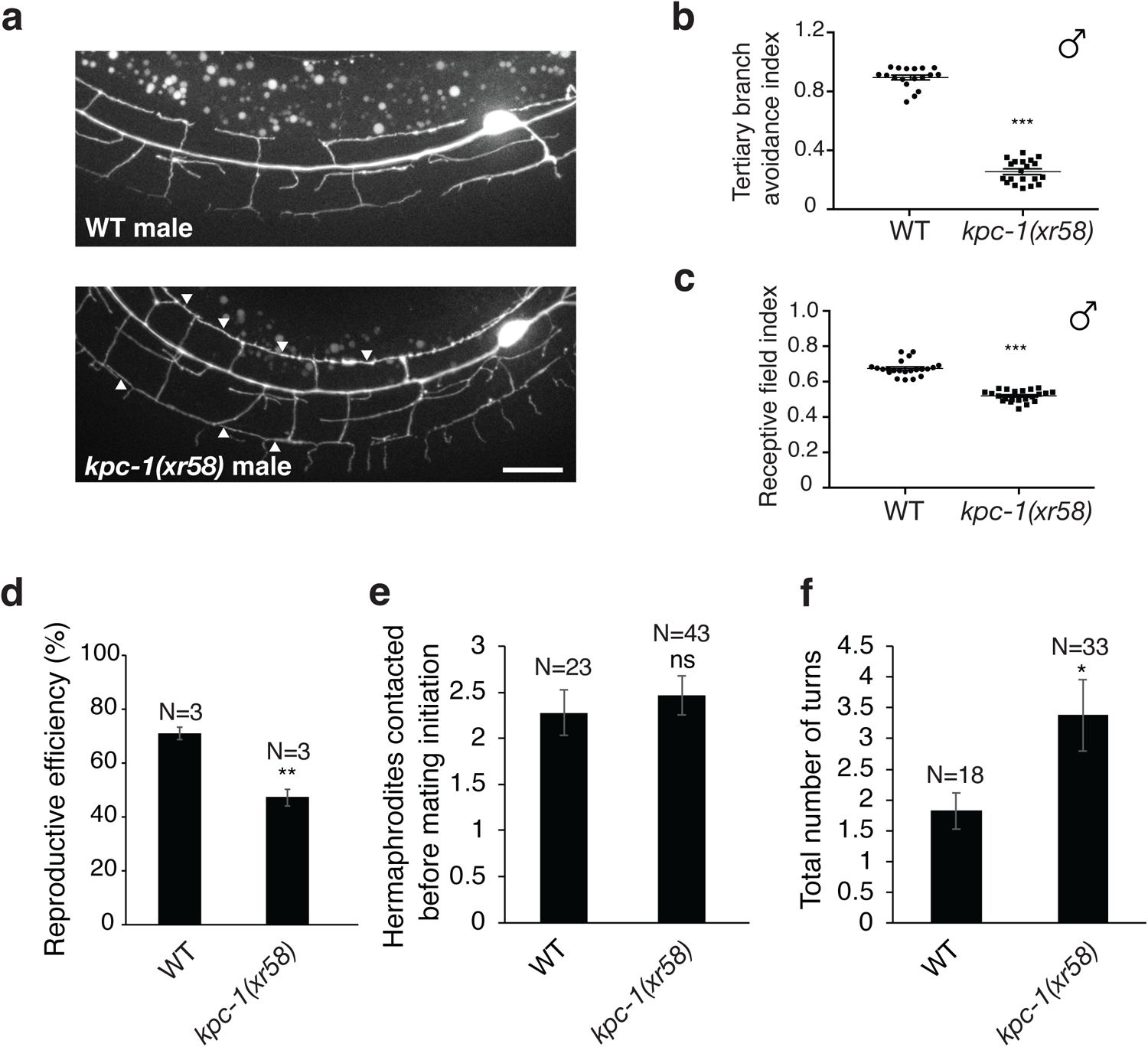
*kpc-1* mutant males exhibit mating defects. (**a**) Images of PVD dendrites in wild-type and *kpc-1(xr58)* mutant males. Arrowheads point to contacts between neighboring tertiary dendrites. Scale bars, 20 μm. (**b**) Tertiary branch avoidance index of the wild-type and *kpc-1(xr58)* mutant males. ***p < 0.001 by a Student’s *t*-test. (**c**) Receptive field index of the wild-type and *kpc-1(xr58)* mutant males. Receptive field index was defined by the ratio of the skin area innervated by the dendrite arbors normalized to the skin area of the animal body, measured from the PVD soma to the end of the mouth (anterior most point in the animal). ***p < 0.001 by a Student’s *t*-test. (**d**) Reproductive efficiencies of wild-type and *kpc-1(xr58)* mutant males. Three repeats were performed for each strain. **p < 0.01 by a Student’s *t*-test. (**e**) The number of hermaphrodites contacted by males before mating initiation. p = 0.86 by a Student’s *t*-test. (**f**) The number of turns the males made around hermaphrodites’ head and tail before locating the vulva. **p < 0.01 by a Student’s *t*-test. Error bars, SEM.

## Discussion

Our results support a model where locally synthesized KPC-1 at branching points and contact points between sibling dendrites down-regulates DMA-1 receptors to promote dendrite branching and self-avoidance (Fig. 7k). A recent paper showed that KPC-1 targeted DMA-1 to endosomes for degradation in PVD neurons(*4*). It is likely that a similar mechanism is utilized in a site-specific manner for secondary dendrite branching(*4*) and tertiary dendrite self-avoidance. KPC-1 encodes a proprotein convertase subtilisin/kexin (PCSK). At a functional level, there are two types of mammalian PCSKs. PCSK3/furin activates substrates by cleavage of precursor proteins, such as TGF-β, whereas PCSK9 inactivates substrates by induced degradation of proteins, such as LDL receptors(*43–46*). Our results indicate that KPC-1 is more closely related to PCSK9 at a functional level in its inactivation of DMA-1. It has been shown that PCSK9 is expressed in the vertebrate CNS, including the cerebellum, and is important for CNS development(*47*). However, it remains to be seen whether PCSK9 plays a similar role to KPC-1 in dendrite branching and self-avoidance by down-regulating dendrite receptors.

It is necessary that the level of DMA-1 receptors on tertiary dendrites is tightly controlled during dendrite outgrowth and self-avoidance (Fig. 7k). For example, the DMA-1 receptor level should be high on tertiary dendrites during dendrite outgrowth. However, the DMA-1 receptor level needs to be down-regulated upon contact between sibling tertiary dendrites, resulting in dendrite retraction (self-avoidance). An important question that remains to be addressed in the future is how the KPC-1 activity is dynamically controlled to regulate corresponding DMA-1 levels. There are two possible regulatory mechanisms that could be responsible: expression versus activation of KPC-1 at the contact points between sibling dendrites. Our observation of the enrichment of *kpc-1* transcripts at the contact points appears to support the former case (Fig. 3e). However, it is not known how *kpc-1* transcripts are targeted to contact points and whether contact-triggered RNA transport is responsible.

Although our results demonstrate the important role of the *kpc-1* 3’UTR in promoting *kpc-1* mRNA transport to the PVD higher order dendrites, questions remain on how the 3’UTR elements interact with the RNA transport system. It was recently reported that 12 conserved RNA-binding protein (RBP)-encoding genes are involved in PVD dendrite development in *C. elegans*(*48*). Among them, loss of function in *cgh-1* and *cpb-3* resulted in an increase in the number of tertiary dendrite branches. The phenotype of increased number of tertiary dendrite branches is also shared by *kpc-1(xr58)* mutants (Fig. 1c). Like *kpc-1*, *cgh-1* and *cpb-3* act cell-autonomously in PVD neurons to control dendrite development. Also, like *kpc-1* transcripts, CGH-1 and CPB-3 proteins are enriched in PVD dendrites, raising the possibility that CGH-1 and CPB-3 may regulate the *kpc-1* function in dendrites by binding to the 3’UTR of *kpc-1* mRNAs, a possibility awaiting to be explored. A Schizophrenia-associated genetic variant in the 3’UTR of the human furin gene, a *kpc-1* homologous gene, was recently shown to result in downregulation of furin expression by acquiring a miR-338-3p binding site, leading to reduced BDNF production. It remains to be seen whether this human furin 3’UTR genetic variant disrupts transport and local translation of the furin mRNA in dendrites and whether it affects dendrite patterning in Schizophrenia-causing neurons(*1*, *49*).

### Experimental Procedures

#### Strains and plasmids

*C. elegans* strains were cultured using standard methods(*50*). All strains were grown at 20°C. Standard protocol was used for the plasmid constructions. Strains and plasmids are listed in Supplementary Tables S1 and S2.

#### Isolation of *kpc-1* alleles from the genetic screen

A forward genetic screen was conducted as previously described. Wild type worms with *xrIs37* marker were treated with EMS. F1 progenies were transferred to single plates, and F2 progenies were screened for profoundly affected dendrite self-avoidance defect. Using a Zeiss fluorescence dissecting microscope. The *kpc-1(xr47)*, *kpc-1(xr58)*, and *kpc-1(xr60)* mutations were identified from a screen of 9,500 genomes.

#### Transgenic animals

Germline transformation of *C. elegans* was performed using standard techniques(*51*). For example, the *Pkpc-1(5K)::kpc-1* transgene was injected at 10 ng/ml along with the coinjection marker *ofm-1::rfp* at 50 ng/μl. Transgenic lines were maintained by following the *ofm-1::rfp* fluorescence.

#### Tertiary branch avoidance index (TBAI)

TBAI was calculated by dividing the number of visible gaps between tertiary branches by the number of menorahs. A menorah-like structure is composed of a collection of 2° (“stem”), 3° (“base”) and 4° (“candles”) branches. In the case where discrete menorahs cannot be easily differentiated due to self-avoidance defects, we counted each menorah as a single 2° (“stem”) branch that extended 3° (“base”) branches.

#### Scatter plots of quaternary dendrite termini

To map out quaternary dendrite termini, we used a set of customized MatLab scripts (MatLAB, Mathworks, Natick, MA) to effectively unroll the worm’s cylindrical surface, quantifying anterior-posterior distances using the coordinate parallel to the body centerline and dorsal-ventral distances along the cylindrical surface. In order to consolidate data from different worms, all scatter plots are scaled to each worm’s circumference. In each scatter plot, the top line indicates the dorsal nerve cord, the bottom line indicates the ventral nerve cord, and the wild-type morphology of the PVD primary dendrite and axon is drawn in green. The distance between the top and bottom lines corresponds to 1/2 of total worm circumference, and the horizontal axis shows a portion of body length equivalent to approximately 3 circumferences.

#### Molecular cloning

All constructs generated for this study are summarized in Table S2. Sequence variants of the *kpc-1* 3’UTR were verified by sequencing.

#### CRISPR-Cas-9 genome editing

To generate deletion of the SLS2 region in the *kpc-1* 3’UTR, we used the Co-CRISPR *dpy-10(cn64)* as a screening method(*52*). Worms were injected with Cas9_sgRNA plasmid and single-stranded oligonucleotide donor (IDT). Both *xr82* and *xr84* SLS2 deletion alleles were confirmed by sequencing.

#### Laser dendritomy

For the femtosecond laser surgery, we used a cavity-dumped Ti:sapphire laser oscillator(*53*, *54*) to generate laser pulses ∼100 fs in duration and 200 kHz in repetition rate. The laser pulses were tightly-focused onto targeted primary dendrites using a Nikon 100x, 1.4 NA oil-immersion objective. The vaporization threshold corresponds to pulse energies of 5-15 nJ. Successful laser dendritomy was confirmed by visualizing the targeted area immediately after surgery. Dendrites were cut at the primary branch 50 μm anterior to the cell body in the early L4 stage when tertiary dendrites undergo self-avoidance. 1°, 2°, 3°, and 4° dendrites in the anterior dendrite arbor were disconnected from the cell body after primary dendritomy. Newly regenerated dendrites can be easily differentiated from the old dendrites 24 hr after surgery due to difference in the fluorescence intensity of the *Pser-2::GFP* dendrite marker.

#### Imaging and quantifying dendrite arbors

The morphology of neuronal cell bodies and dendrites was based on high-magnification Z-stacks using a Zeiss 60x, 1.4 NA oil-immersion objective. For laser dendritomy, we mounted individual animals on 2% agar pads and anaesthetized them with 3 mM sodium azide, the lowest possible concentration to keep adult animals immobilized. Laser dendritomy was performed and worms were recovered within 10 minutes of sodium azide treatment. Recovered worms were placed on fresh plates with bacterial foods and imaged 24 hours after dendritomy using a Hamamatsu ORCA AG camera. For imaging and quantifying uninjured dendrite arbors, young adult animals were mounted on 2% agar pads and anaesthetized with 20 mM sodium azide.

The dendrite length of regenerating neurons was quantified 24 hours after surgery. Dendrite lengths were calculated as the actual contour length between the injury site and dendrite termini measured along the cylindrical surface of each worm, by tracing dendrites through a 3-dimensional image stack. P values for the length measurements were calculated using a student’s t-Test.

#### Fluorescence microscopy

Worms were mounted on 2% agarose pad and anaesthetized with 20 mM sodium azide solution. Images were acquired using a 40x, 1.4 NA objective on a Zeiss Axio Imager M2 microscope with a Hamamatsu ORCA-Flash4.0 LT+ camera. Images were processed by z-stack projections.

#### Fluorescence recovery after photoconversion

Worms were mounted on 2% agarose pad and anaesthetized with 7.5 mM tetramisole. Photoconversion of Kaede was performed under 405 nm laser on the region of interest (ROI). Fluorescence quantification of ROI intensity was measured every 5 seconds over 6 minutes following photoconversion (Zeiss Imaging Browser). At each time point, percentage green fluorescence signal recovery was determined by (F_n_-F_0_)/(F_p_-F_0_), where F_n_ = fluorescence intensity n seconds after photoconversion; F_0_ = fluorescence intensity immediately after photoconversion; F_p_ = fluorescence intensity right before photoconversion. All images were taken in live animals using a 40x, 1.3 NA objective and 488 nm and 561 nm laser on a Zeiss LSM 880 confocal microscope.

#### Male mating behavior assays

Male mating behavioral assays were conducted using standard methods(*55*). Reproductive efficiencies were conducted by placing six males with seven hermaphrodite carrying *unc-36(e251)* recessive marker and results were calculated by the number of cross-progeny over total number progeny. Three repeats were conducted on separate days. Number of hermaphrodites contacted before mating initiation were counted within 5 minutes of male first contacted the hermaphrodites. Total number of turns around hermaphrodites during mating were counted before males successfully identify the vulva.

#### Statistics

Average data of dendrite number, dendrite length, TBAI, receptive field index, length ratio of receptive field, and reporter expression intensity are presented as means ± SEM. Data of % PVD neurons with secondary dendrite branching defects, defective dendrite self-avoidance, and dendritic signals are presented as proportions ± SEP. Statistical analyses were carried out by Student’s *t*-tests, two-proportion *Z*-tests, or one-way ANOVA with Tukey’s or Dunnett’s tests using GraphPad Prism 7.0 or the Primer of Biostatistics software.

## Author Contributions

M.S. conceived, designed, performed, and analyzed experiments, made constructs, and drafted the article related to figures 4, 5, 8, and S3-S5. Y.Z. conceived, designed, performed, and analyzed experiments, and made constructs. T.F. performed genetic screens, identified dendrite self-avoidance mutants, determined receptive field index and length ratio of receptive field in PVD dendrite arbors, and drew scatter plots of 4° dendrite termini using MATLAB. N.Z. imaged and quantified green fluorescent MS2 capsid proteins in PVD dendrites. E.K. performed male mating behavioral assays and analyzed results. C.F.C. conceived, designed, and analyzed experiments. C.C. conceived, designed, analyzed and interpreted data, and drafted the article.

## Acknowledgments

This work was funded by grants from the March of Dimes Foundation (C.C.), the Whitehall Foundation Research Award (C.C.), the National Science Foundation (IOS-1455758 to C.C.), and the National Institute of General Medical Sciences of the National Institutes of Health (R01GM111320 to C.C.). We thank Oliver Hobert lab for Whole Genome Sequencing, Daniel J. Dickinson and Bob Goldstein for providing reagents and CRISPR protocols, Kana Hamada for confocal imaging protocols, Ryan Weihsiang Lin for imaging analysis, Seema Sheoran for unpublished observation, Evguenia Ivakhnitskaia for critical reading the manuscript, Hui Chiu for critical reading the manuscript, providing critical initial data, and making main figures, Kang Shen for the *dma-1(wy1246) [dma-1::GFP]* allele, the *Caenorhabditis* Genetics Center for the *kpc-1(gk8)* strain, and the WormBase for readily accessible information.

