## Supplemental file for "The *kpc-1* 3’UTR facilitates dendritic transport and translation of mRNAs for dendrite arborization of a mechanosensory neuron important for male courtship"

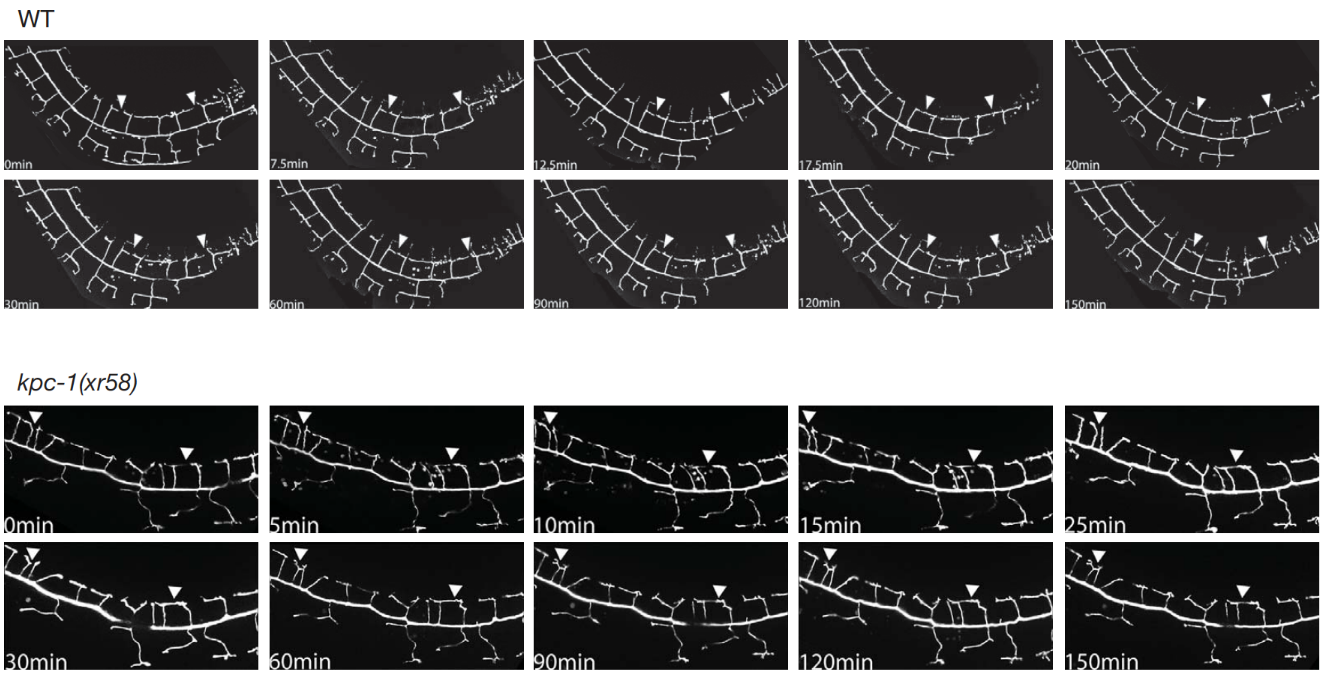
WT
0min
7.5min
12.5min
17.5min
20min
30min
60min
90min
120min
150min
kpc-1(xr58)
0min
5min
10min
15min
25min
30min
60min
90min
120min
150min

**Supplementary Figure 1. Time-lapse imaging of PVD dendritic arborization in wild type and *kpc-1(xr58)* mutants.** Time series images of tertiary dendritic arborization in wild type and *kpc-1(xr58)* mutants are shown. PVD neurons were imaged at 2.5-30 min intervals for 2.5 hours at the late L3 stage using confocal fluorescence microscopy. Arrowheads point to the dynamic contact points.

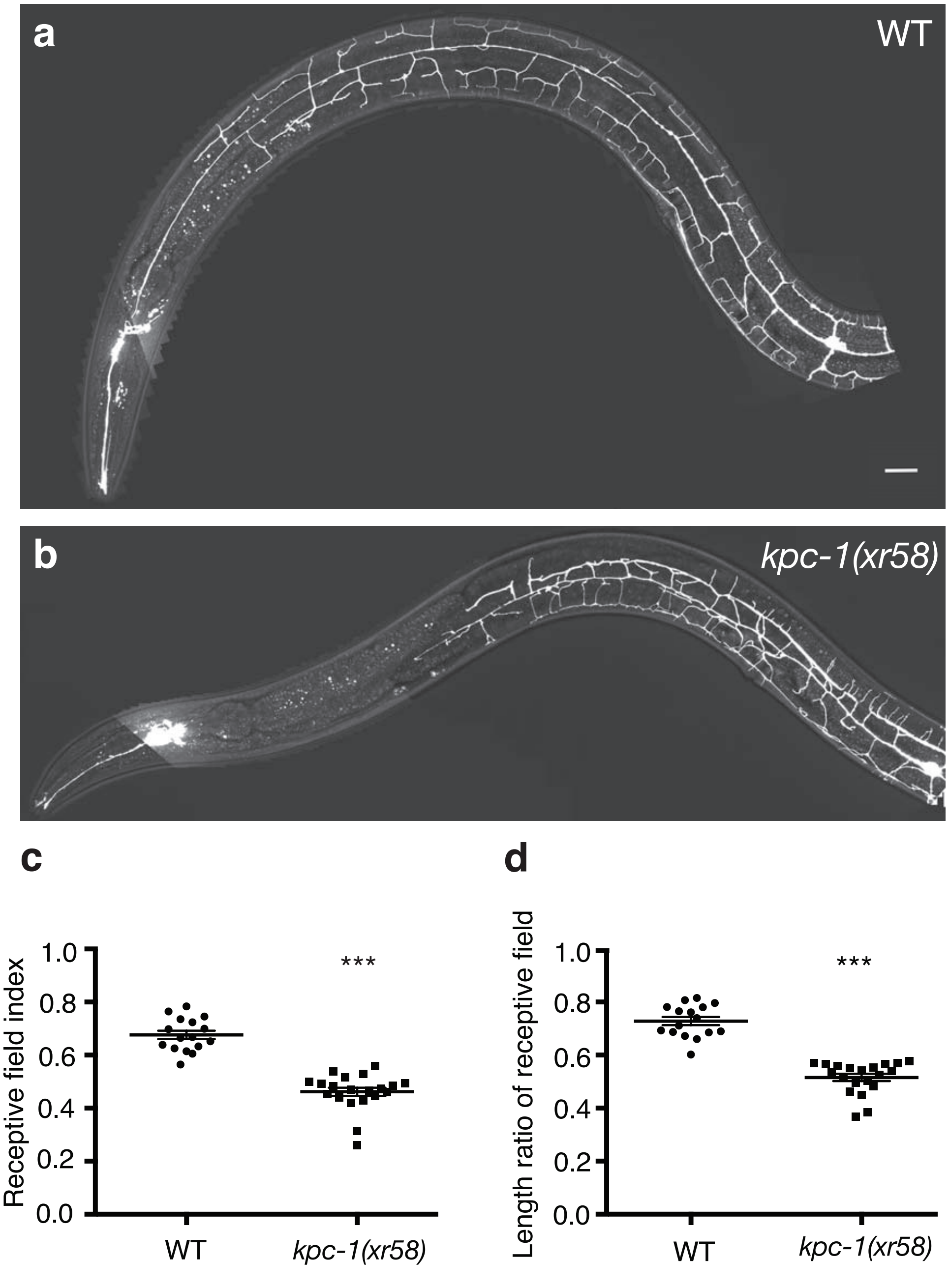


**Supplementary Figure 2**

**Supplementary Figure 2. Measurement of sensory coverage of skin by PVD dendrite arbors.** (**a, b**) Superimposed images of PVD dendrite arbors and animal bodies in wild type (**a**) and *kpc-1(xr58)* mutants (**b**). (**c**) Receptive field index was defined by the ratio of the skin area innervated by the dendrite arbors normalized to the skin area of the animal body, measured from the PVD soma to the end of the mouth (anterior most point in the animal). (**d**) Length ratio of receptive field was determined by the length of the PVD dendrite arbors along the anterior-posterior axis normalized to the distance between the PVD soma and the end of the mouth along the anterior-posterior axis.

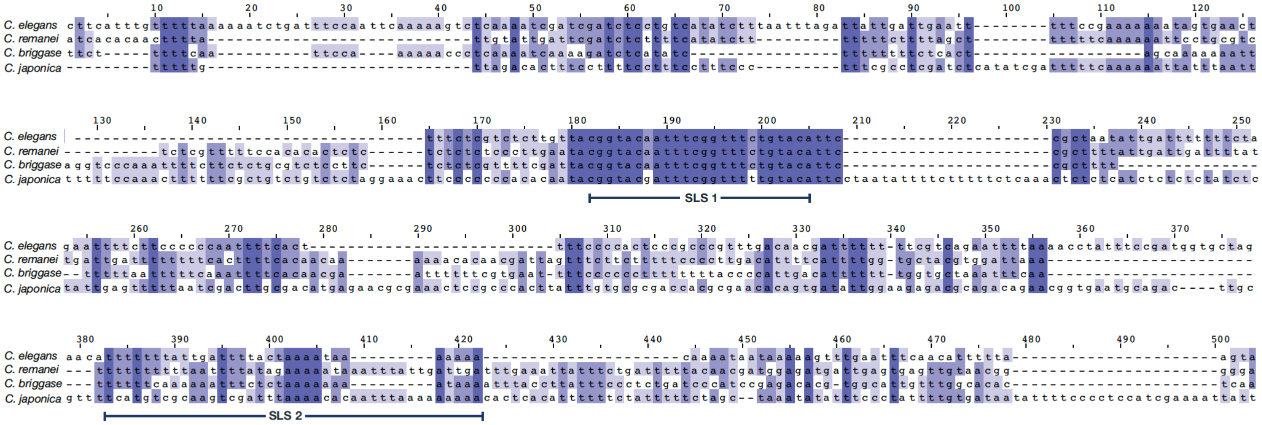
C. elegans
C. remanei
C. briggase
C. japonica
SLS 1
SLS 2

**Supplementary Figure 3. Alignment of *kpc-1* 3'UTR sequences from *Caenorhabditis* species.** MAFFT alignment of 4 *kpc-1 3'UTR* sequences shown here corresponds to the following subsequences: *C. elegans*, nt 1-357; *C. remanei*, nt 1-366; *C. briggsae*, nt 1-340; *C. japonica*, nt 1-452. Stem-loop structure 1 (SLS1) and 2 (SLS2) regions are indicated.

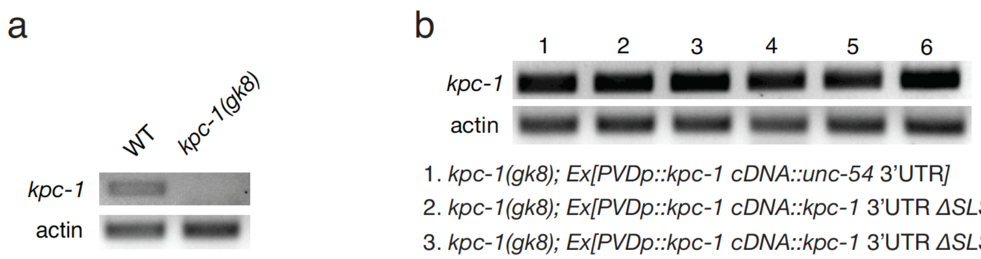


1. *kpc-1(gk8); Ex[PVDp::kpc-1 cDNA::unc-54* 3'UTR*]*
2. *kpc-1(gk8); Ex[PVDp::kpc-1 cDNA::kpc-1* 3'UTR *ΔSLS1]*
3. *kpc-1(gk8); Ex[PVDp::kpc-1 cDNA::kpc-1* 3'UTR *ΔSLS2]*
4. *kpc-1(gk8); Ex[PVDp::kpc-1 cDNA::kpc-1* 3'UTR *SLS2 mutated base pairing disrupted]*
5. *kpc-1(gk8); Ex[PVDp::kpc-1 cDNA::kpc-1* 3'UTR*]*
6. *kpc-1(gk8); Ex[PVDp::kpc-1 cDNA::kpc-1* 3'UTR *SLS2 mutated base pairing preserved]*

**Supplementary Figure 4. The *kpc-1* mRNA expression levels are comparable among different *kpc-1* transgenes. (a)** RT-PCR was performed to determine the endogenous *kpc-1* mRNA levels in wild-type and *kpc-1(gk8)* mutants using an oligo(dT) RT primer and a pair of specific PCR primers to amplify a *kpc-1* fragment that is uncovered in the *kpc-1(gk8)* deletion allele. **(b)** The transgenic *kpc-1* mRNA levels in *kpc-1(gk8)* mutants expressing different *kpc-1* transgenes measured by RT-PCR. Actin was used as an internal control.

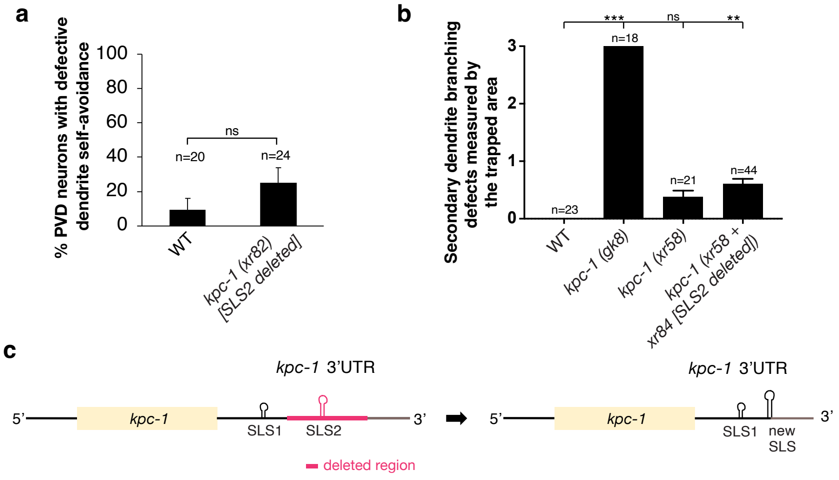
a
% PVD neurons with defective dendrite self-avoidance
100
80
60
40
20
0
ns
n=20
n=24
WT
kpc-1 (xr82) [SLS2 deleted]
b
Secondary dendrite branching defects measured by the trapped area
3
2
1
0
***
ns
**
n=18
n=21
n=44
n=23
WT
kpc-1 (gk8)
kpc-1 (xr58)
kpc-1 (xr58 + xr84 [SLS2 deleted])
c
kpc-1 3'UTR
5'
kpc-1
SLS1
SLS2
3'
deleted region
kpc-1 3'UTR
5'
kpc-1
SLS1
new SLS
3'

**Supplementary Figure 5. CRISPR engineering of *kpc-1* 3'UTR and its effects on PVD dendrite arborization. (a)** Percentage of PVD neurons with defective dendrite self-avoidance. *kpc-1(xr82) [SLS2 deleted]* was created by the CRISPR engineering. Defective dendrite self-avoidance was defined as more than one tertiary dendrite contact point for each PVD neuron. **(b)** Secondary dendrite branching defects measured by the trapped area. *kpc-1(xr58 + xr84[SLS2 deleted])* was generated in the *xr58* background in a separate CRISPR event from *xr82*, and both *xr82* and *xr84* alleles contain the same deletion that uncovers SLS2. The defect of secondary dendrite branching was qualitatively determined to be no (0), mild (1), moderate (2), and severe (3) defects. **$p<0.01$ and ***$p<0.001$ by one-way ANOVA with Dunnett's test. **(c)** Schematic of the new SLS generated following the deletion in *xr82* and *xr84* alleles.

**<u>Supplementary Table 1</u>. A strain list**

| Strain | Mutations | Integrated transgenes | Extrachromosomal transgenes |
|---|---|---|---|
| XN1467 | | *xrIs37[PF49H12.4::GFP; Pmec-3::mcherry] IV* | |
| XN1764 | *kpc-1(xr58) I* | *xrIs37 IV* | |
| XN1820 | *kpc-1(gk8) I* | *xrIs37 IV* | |
| XN1850 | *kpc-1(xr58) I* | *xrIs37 IV* | *xrEx617[Pkpc-1(5K)::kpc-1::kpc-1 3'UTR(3k)* 10ng/µl*; ofm-1::rfp]* |
| XN1851 | *kpc-1(xr58) I* | *xrIs37 IV* | *xrEx618[Pkpc-1(5K)::kpc-1::kpc-1 3'UTR(3k)* 10ng/µl*; ofm-1::rfp]* |
| XN1848 | *kpc-1(gk8) I* | *xrIs37 IV* | *xrEx615[Pkpc-1(5K)::kpc-1::kpc-1 3'UTR(3k)* 10ng/µl*; ofm-1::rfp]* |
| XN1849 | *kpc-1(gk8) I* | *xrIs37 IV* | *xrEx616[Pkpc-1(5K)::kpc-1::kpc-1 3'UTR(3k)* 10ng/µl*; ofm-1::rfp]* |
| XN1872 | *kpc-1(xr58) I* | *xrIs37 IV* | *xrEx628[Pkpc-1(5K)::kpc-1s::kpc-1 3'UTR(3k)* 10ng/µl*; ofm-1::rfp]* |
| XN1873 | *kpc-1(xr58) I* | *xrIs37 IV* | *xrEx629[Pkpc-1(5K)::kpc-1s::kpc-1 3'UTR(3k)* 10ng/µl*; ofm-1::rfp]* |
| XN1875 | *kpc-1(gk8) I* | *xrIs37 IV* | *xrEx631[Pkpc-1(5K)::kpc-1s::kpc-1 3'UTR(3k)* 10ng/µl*; ofm-1::rfp]* |
| XN1876 | *kpc-1(gk8) I* | *xrIs37 IV* | *xrEx632[Pkpc-1(5K)::kpc-1s::kpc-1 3'UTR(3k)* 10ng/µl*; ofm-1::rfp]* |
| XN1879 | *kpc-1(xr58) I* | *xrIs37 IV* | *xrEx635[PF49H12.4::kpc-1s::kpc-1 3'UTR(3k)* 10ng/µl*; ofm-1::rfp]* |
| XN1880 | *kpc-1(xr58) I* | *xrIs37 IV* | *xrEx636[PF49H12.4::kpc-1s::kpc-1 3'UTR(3k)* 10ng/µl*; ofm-1::rfp]* |
| XN1877 | *kpc-1(gk8) I* | *xrIs37 IV* | *xrEx633[PF49H12.4::kpc-1s::kpc-1 3'UTR(3k)* 10ng/µl*; ofm-1::rfp]* |
| XN1878 | *kpc-1(gk8) I* | *xrIs37 IV* | *xrEx634[PF49H12.4::kpc-1s::kpc-1 3'UTR(3k)* 10ng/µl*; ofm-1::rfp]* |
| XN1822 | *kpc-1(xr58) I* | *xrIs37 IV* | *xrEx596[PF49H12.4::kpc-1s::unc-54 3'UTR* 25ng/µl*; ofm-1::rfp]* |
| XN1824 | *kpc-1(xr58) I* | *xrIs37 IV* | *xrEx598[PF49H12.4::kpc-1s::unc-54 3'UTR* 25ng/µl*; ofm-1::rfp]* |
| XN1829 | *kpc-1(gk8) I* | *xrIs37 IV* | *xrEx596[PF49H12.4::kpc-1s::unc-54 3'UTR* 25ng/µl*; ofm-1::rfp]* |
| XN1830 | *kpc-1(gk8) I* | *xrIs37 IV* | *xrEx598[PF49H12.4::kpc-1s::unc-54 3'UTR* 25ng/µl*; ofm-1::rfp]* |
| XN1836 | *kpc-1(xr58) I* | *xrIs37 IV* | *xrEx602[PF49H12.4::kpc-1L::unc-54 3'UTR* 25ng/µl*; ofm-1::rfp]* |
| XN1837 | *kpc-1(xr58) I* | *xrIs37 IV* | *xrEx603[PF49H12.4::kpc-1L::unc-54 3'UTR* 25ng/µl*; ofm-1::rfp]* |
| XN1838 | *kpc-1(gk8) I* | *xrIs37 IV* | *xrEx605[PF49H12.4::kpc-1L::unc-54 3'UTR* 25ng/µl*; ofm-1::rfp]* |
| XN1839 | *kpc-1(gk8) I* | *xrIs37 IV* | *xrEx606[PF49H12.4::kpc-1L::unc-54 3'UTR* 25ng/µl*; ofm-1::rfp]* |
| XN1840 | *kpc-1(xr58) I* | *xrIs37 IV* | *xrEx607[PF49H12.4::kpc-1s::unc-54 3'UTR* 100ng/µl*; ofm-1::rfp]* |
| XN1841 | *kpc-1(xr58) I* | *xrIs37 IV* | *xrEx608[PF49H12.4::kpc-1s::unc-54 3'UTR* 100ng/µl*; ofm-1::rfp]* |
| XN1842 | *kpc-1(gk8)I* | *xrIs37 IV* | *xrEx609[PF49H12.4::kpc-1s::unc-54 3'UTR* 100ng/µl*; ofm-1::rfp]* |
| XN1843 | *kpc-1(gk8) I* | *xrIs37 IV* | *xrEx610[PF49H12.4::kpc-1s::unc-54 3'UTR* 100ng/µl*; ofm-1::rfp]* |
| XN1901 | | | *xrEx654[Pkpc-1(5K)::GFP::kpc-1 3'UTR(3k); PF49H12.4::mcherry; odr-1::rfp]* |
| XN1902 | | | *xrEx655[Pkpc-1(5K)::GFP::kpc-1 3'UTR(3k); PF49H12.4::mcherry; odr-1::rfp]* |
| XN1856 | | *xrIs37 IV* | *xrEx622[Pkpc-1(5K)::kpc-1s* 10ng/µl*; ofm-1::rfp]* |
| XN1857 | | *xrIs37 IV* | *xrEx624[Pkpc-1(5K)::kpc-1s* 10ng/µl*; ofm-1::rfp]* |
| XN1949 | | *xrIs37 IV* | *wyEx4725[Pser-2::dma-1* 80ng/µl*]* |
| XN1946 | *kpc-1(xr58) I* | *xrIs37 IV* | *wyEx4725[Pser-2::dma-1* 80ng/µl*]* |
| XN1950 | | *xrIs37 IV* | *xrEx686[Pkpc-1(5K)::kpc-1s* 10ng/µl*; ofm-1::rfp]; wyEx4725[ser-2p::dma-1* 80ng/µl*]* |
| XN1951 | | *xrIs37 IV* | *xrEx687[Pkpc-1(5K)::kpc-1s* 10ng/µl*; ofm-1::rfp]; wyEx4725[ser-2p::dma-1* 80ng/µl*]* |
| XN2012 | | | *xrEx719[Pser-2::NLS::MS2::GFP; Podr-1::rfp]* |
| XN2095 | | | *xrEx766[Pser-2::NLS::MS2::GFP; Pser-2::kpc-1s::24xms2 bs::unc-54 3'UTR; Podr-1::rfp]* |
| XN1999 | *kpc-1(xr58) I* | | *xrEx696[Pser-2::dma-1::mcherry::unc-54 3'UTR; egl-20::gfp]* |
| XN1993 | *kpc-1(xr47) I* | | *xrEx696[Pser-2::dma-1::mcherry::unc-54 3'UTR; egl-20::gfp]* |
| XN2389 | | | *xrEx890[Pser-2prom3::NLS::MS2::GFP* 10ng/ul*; Pser-2:kpc-1s::24xMS2bs::kpc-13'UTR* 10ng/µl*; Podr-1::RFP]* |
| XN2390 | | | *xrEx891[Pser-2prom3::NLS::MS2::GFP* 10ng/ul*; Pser-2:kpc-1s::24xMS2bs::kpc-13'UTR* 10ng/µl*; Podr-1::RFP]* |
| XN2481 | *kpc-1(gk8) I* | *xrIs37 IV* | *xrEx955[PF49H12.4::kpc-1s::(SLS1 deleted) kpc-1 3'UTR* 10ng/µl*;Podr-1::RFP* 50ng/µl*]* |
| XN2482 | *kpc-1(gk8) I* | *xrIs37 IV* | *xrEx956[PF49H12.4::kpc-1s::(SLS1 deleted) kpc-1 3'UTR* 10ng/µl*;Podr-1::RFP* 50ng/µl*]* |
| XN2483 | *kpc-1(gk8) I* | *xrIs37 IV* | *xrEx957[PF49H12.4::kpc-1s::(SLS1 deleted) kpc-1 3'UTR* 10ng/µl*;Podr-1::RFP* 50ng/µl*]* |
| XN2520 | *kpc-1(gk8) I* | *xrIs37 IV* | *xrEx978[PF49H12.4::kpc-1s::(SLS2 deleted) kpc-1 3'UTR* 10ng/µl*;Podr-1::RFP* 50ng/µl*]* |
| XN2521 | *kpc-1(gk8) I* | *xrIs37 IV* | *xrEx979[PF49H12.4::kpc-1s::(SLS2 deleted) kpc-1 3'UTR* 10ng/µl*;Podr-1::RFP* 50ng/µl*]* |
| XN2522 | *kpc-1(gk8) I* | *xrIs37 IV* | *xrEx980[PF49H12.4::kpc-1s::(SLS2 deleted) kpc-1 3'UTR* 10ng/µl*;Podr-1::RFP* 50ng/µl*]* |
| XN2550 | *kpc-1(gk8) I* | *xrIs37 IV* | *xrEx1004[PF49H12.4::kpc-1s::(SLS2 deleted) kpc-1 3'UTR* 10ng/µl*;Podr-1::RFP* 50ng/µl*]* |
| XN2551 | *kpc-1(gk8) I* | *xrIs37 IV* | *xrEx1005[PF49H12.4::kpc-1s::(SLS2 deleted) kpc-1 3'UTR* 10ng/µl*;Podr-1::RFP* 50ng/µl*]* |
| XN2552 | *kpc-1(gk8) I* | *xrIs37 IV* | *xrEx1006[PF49H12.4::kpc-1s::(SLS2 deleted) kpc-1 3'UTR* 10ng/µl*;Podr-1::RFP* 50ng/µl*]* |
| XN2570 | *kpc-1(gk8) I* | *xrIs37 IV* | *xrEx1020[PF49H12.4::kpc-1s::(SLS2 mutated base pair disrupted) kpc-1 3'UTR* 10ng/µl*; Podr-1::RFP* 50ng/µl*]* |
| XN2571 | *kpc-1(gk8) I* | *xrIs37 IV* | *xrEx1021[PF49H12.4::kpc-1s::(SLS2 mutated base pair disrupted) kpc-1 3'UTR* 10ng/µl*; Podr-1::RFP* 50ng/µl*]* |
| XN2572 | *kpc-1(gk8) I* | *xrIs37 IV* | *xrEx1022[PF49H12.4::kpc-1s::(SLS2 mutated base pair disrupted) kpc-1 3'UTR* 10ng/µl*; Podr-1::RFP* 50ng/µl*]* |
| XN2581 | *kpc-1(gk8) I* | *xrIs37IV* | *xrEx1041[PF49H12.4::kpc-1s::(SLS2 mutated base pair preserved) kpc-1 3'UTR* 10ng/µl*; Podr-1::RFP* 50ng/µl*]* |
| XN2582 | *kpc-1(gk8) I* | *xrIs37IV* | *xrEx1042[PF49H12.4::kpc-1s::(SLS2 mutated base pair preserved) kpc-1 3'UTR* 10ng/µl*; Podr-1::RFP* 50ng/µl*]* |
| XN2654 | | | *xrEx1071[Pser-2prom3::kaede::kpc-1 3'UTR* 50ng/µl*; Podr-1::RFP* 50ng/µl*]* |
| XN2655 | | | *xrEx1072[Pser-2prom3::kaede::kpc-1 3'UTR* 50ng/µl*; Podr-1::RFP* 50ng/µl*]* |
| XN2656 | | | *xrEx1073[Pser-2prom3::kaede::kpc-1 3'UTR* 50ng/µl*; Podr-1::RFP* 50ng/µl*]* |
| XN2657 | | | *xrEx1074[Pser-2prom3::kaede::kpc-1 3'UTR* 50ng/µl*; Podr-1::RFP* 50ng/µl*]* |
| XN2658 | | | *xrEx1075[Pser-2prom3::kaede::kpc-1 3'UTR* 50ng/µl*; Podr-1::RFP* 50ng/µl*]* |
| XN2659 | | | *xrEx1076[Pser-2prom3::kaede::kpc-1 3'UTR* 50ng/µl*; Podr-1::RFP* 50ng/µl*]* |
| XN2927 | *kpc-1(xr82) I* | | |
| XN2939 | *kpc-1(xr58xr84) I* | | |

| | | | |
|---|---|---|---|
| XN2955 | *kpc-1(xr47) I* | *dma-1(wy1246) I* | |
| XN2956 | *kpc-1(xr58) I* | *dma-1(wy1246) I* | |
| TV24913 | | *dma-1(wy1246)* [DMA-1::GFP] *I* | |
| CHB392 | | | *hmnEx113[pCY2(ser-2prom3::kaede)]* |
| XN2790 | *rps-18(ok3353) IV* | *juSi94 [GFP11::rps-18 + Cbr-unc-119(+)] II. juIs409 [rgef-1p::GFP1-10 + ttx-3p::RFP] IV* | *xrEx1146[Pser2::mCherry* 10ng/µl*; Podr-1::RFP* 50ng/µl*]* |
| CB251 | *unc-36(e251) III* | | |

**<u>Supplementary Table 2</u>. A plasmid list**

| Plasmids |
|---|
| *Pkpc-1(5kb)::kpc-1::kpc-1*$^{3'UTR}$*(3kb)* was a PCR-amplified genomic fragment. |
| *PF49H12.4::kpc-1s::unc-54*$^{3'UTR}$ was constructed by RT-PCR amplification of the *kpc-1s* cDNA, followed by digestion with NheI and EcoRI and ligation to the *PF49H12.4::unc-54*$^{3'UTR}$*-pSM* vector. |
| *PF49H12.4::kpc-1l::unc-54*$^{3'UTR}$ was constructed by RT-PCR amplification of the *kpc-1l* cDNA, followed by digestion with NheI and EcoRI and ligation to the *PF49H12.4::unc-54*$^{3'UTR}$*-pSM* vector. |
| *Pdpy-7::kpc-1s::unc-54* $^{3'UTR}$ was constructed by PCR amplification of a 0.4-kb *dpy-7* promoter fragment, followed by digestion with FseI and NheI and ligation to the *kpc-1s::unc-54*$^{3'UTR}$*-pSM* vector. |
| *PF49H12.4::kpc-1s::kpc-1*$^{3'UTR}$*(3kb)* was constructed by PCR amplification of a 3-kb *kpc-1*$^{3'UTR}$ genomic fragment, followed by digestion with EcoRI (blunted) and SpeI and ligation to the *PF49H12.4::kpc-1s-pSM* vector. |
| *Pkpc-1(5kb)::kpc-1s::kpc-1*$^{3'UTR}$*(3kb)* was constructed by PCR amplification of a 5-kb *kpc-1* promoter fragment, followed by digestion with FseI and AscI and ligation to the *kpc-1s::kpc-1*$^{3'UTR}$*(3kb)-pSM* vector. |
| *Pkpc-1(5kb)::GFP::kpc-1*$^{3'UTR}$*(3kb)* was constructed by PCR amplification of a 5-kb *kpc-1* promoter fragment, followed by digestion with FseI and AscI and ligation to the *GFP::kpc-1*$^{3'UTR}$*(3kb)-pSM* vector. |
| *PF49H12.4::kpc-1s::GFP::kpc-1*$^{3'UTR}$*(3kb)* was constructed by PCR amplification of the *kpc-1s* cDNA, followed by digestion with AscI and XhoI and ligation to the *PF49H12.4::GFP::kpc-1*$^{3'UTR}$*(3kb)-pSM* vector. |
| *Pser-2::kpc-1s::GFP::kpc-1*$^{3'UTR}$*(3kb)* was constructed by PCR amplification of a 4-kb *ser-2* promoter fragment, followed by digestion with FseI and AscI and ligation to the *kpc-1s::GFP::kpc-1*$^{3'UTR}$*(3kb)-pSM* vector. |
| *Pser-2::kpc-1s::24x MS2 binding site::kpc-1 3'UTR: kpc-1s* was amplified by PCR from *pSM_Pser-2prom3::kpc-1s::GFP::kpc-1 3'UTR* and digested with NheI and SacI and ligated into *pSM*; 24xMS*2 binding site* was amplified by PCR from *phage_cmv::cfp::24*MS2* and digested with SacI and SpeI and ligated into *pSM_kpc-1s*; *kpc-1 3'UTR* was amplified by PCR from N2 gDNA and digested with XhoI and SbfI and ligated into *pSM_kpc-1s::24x MS2 binding site*; *Pser-2prom3* was digested with FseI and AscI and ligated into *pSM_kpc-1s::24x MS2 binding site::kpc-1 3'UTR* |
| *Pser-2::kpc-1s::24x MS2 binding site::unc-54 3'UTR: unc-54 3'UTR* was digested with SacI (blunted with T4 DNA Polymerase) and SpeI from *pSM* and ligated into *pSM_kpc-1s::24x MS2 binding site* which was digested by SbfI (blunted with T4 DNA Polymerase) and SpeI; *Pser-2prom3* was digested with FseI and AscI from *pSM_Pser-2prom3* and ligated into *pSM_kpc-1s::24x MS2 binding site::unc-54 3'UTR* |
| *PF49H12.4::kpc-1s::kpc-1 3'UTR (SLS1 deleted)* SLS1 deletion was introduced by Q5® Site-Directed Mutagenesis Kit (New England Biolabs) using *PF49H12.4::kpc-1s::kpc-1*$^{3'UTR}$*(3kb)* as template. |
| *PF49H12.4::kpc-1s::kpc-1 3'UTR (SLS2 deleted)* SLS2 deletion was introduced by Q5® Site-Directed Mutagenesis Kit (New England Biolabs) using *PF49H12.4::kpc-1s::kpc-1*$^{3'UTR}$*(3kb)* as template. |
| *PF49H12.4::kpc-1s::kpc-1 3'UTR (SLS2 mutated base pair disrupted)* SLS2 mutated base pair disrupted point mutations was introduced by Q5® Site-Directed Mutagenesis Kit (New England Biolabs) using *PF49H12.4::kpc-1s::kpc-1*$^{3'UTR}$*(3kb)* as template. |
| *PF49H12.4::kpc-1s::kpc-1 3'UTR (SLS2 mutated base pair preserved)* SLS2 mutated base pair preserved point mutations was introduced by Q5® Site-Directed Mutagenesis Kit (New England Biolabs) using *PF49H12.4::kpc-1s::kpc-1*$^{3'UTR}$*(3kb)* as template. |
| *Pser2p3::kaede::kpc-1 3'UTR* was constructed by PCR amplification of *kpc-1 3'UTR* fragment, followed by digestion with NotI and PspOMI and ligation to the pCY2 *ser-2prom3::kaede* vector. |
